# The LncRNA *Carmn* is a Critical Regulator for Gastrointestinal Smooth Muscle Contractile Function and Motility

**DOI:** 10.1101/2022.06.28.498024

**Authors:** Xiangqin He, Kunzhe Dong, Jian Shen, Guoqing Hu, James D. Mintz, Reem T. Atawia, Juanjuan Zhao, Xiuxu Chen, Robert W. Caldwell, Meixiang Xiang, David W. Stepp, David J. Fulton, Jiliang Zhou

**Affiliations:** Department of Pharmacology & Toxicology, Medical College of Georgia, Augusta University, Augusta, GA 30912; Department of Cardiology, The Second Affiliated Hospital, Zhejiang University School of Medicine, Hangzhou, China. 310009; Vascular Biology Center, Medical College of Georgia, Augusta University, Augusta, GA30912; Department of Pathology and Laboratory Medicine, Loyola University Health System, Maywood, IL 60153; Department of Physiology, Medical College of Georgia, Augusta University, Augusta, GA 30912

**Author notes:** **Corresponding author Jiliang Zhou,** M.D./Ph.D., Department of Pharmacology & Toxicology, Medical College of Georgia, Augusta University, 1459 Laney Walker Blvd, Augusta, GA 30912. **Author contributions to manuscript:** J.Z. conceived and supervised the project. X.H., K.D. and J.S. designed and performed experiments. X.H. wrote the manuscript. G.H., J.M., R.A., Ju.Z., X.C., R.C., M.X., J.M., D.S. and D.F. performed experiments and/or edited the manuscript. **Data availability statement:** The original data, analytic methods, and materials will be made available to other researchers upon reasonable requests.

**Keywords:** *Carmn*, lncRNA, Smooth muscle, Contractility, Pseudo-obstruction, Visceral myopathy

## Abstract

**Background & aims:** Visceral smooth muscle cells (SMCs) are an integral component of the gastrointestinal (GI) tract and are critical for regulating motility. SMC contraction is regulated by changes in post-translational signaling and the state of differentiation. Impaired SMC contraction is associated with significant morbidity and mortality but the mechanisms regulating the expression levels of SMC-specific contractile proteins, including the role of long non-coding RNAs (lncRNAs), remains largely unexplored. Herein, we have uncovered an important role of *Carmn* (Cardiac mesoderm enhancer-associated noncoding RNA), a SMC-specific lncRNA, in regulating the phenotype of visceral SMCs of the GI tract.

**Methods:** Analysis of GTEx and publicly available single-cell RNA sequencing (scRNA-seq) datasets from embryonic, adult human and mouse GI tissues were used to identify SMC-specific lncRNAs. The functional role of *Carmn* was investigated using a novel GFP knock-in (KI) reporter/knockout (KO) mouse model. Bulk RNA sequencing (RNA-seq) and single nuclei RNA sequencing (snRNA-seq) of colonic muscularis were used to investigate underlying mechanisms.

**Results:** Unbiased in silico analyses and GFP expression patterns in *Carmn* GFP KI mice revealed that *Carmn* is specifically expressed in SMCs in human and mouse GI tract. Premature lethality was observed in global *Carmn* KO (gKO) and inducible SMC-specific KO (iKO) mice due to colonic pseudo-obstruction, severe distension of the GI tract with blockages in cecum and colon segments. Histology, whole-gut GI transit time and muscle myography analysis revealed severe dilation, significantly delayed GI transit and impaired GI contractility in *Carmn* KO mice versus control mice. Bulk RNA-seq of colonic muscularis revealed that *Carmn* deficiency promotes SMC de-differentiation as evidenced by up-regulation of extracellular matrix genes and down-regulation of SMC contractile genes including *Mylk*, a key regulator of SMC contraction. SnRNA-seq further revealed SMC *Carmn* deficiency not only compromised myogenic motility by reducing expression of contractile genes but also impaired neurogenic motility by disrupting cell-cell connectivity in the colonic muscularis. These findings may have translational significance as silencing *CARMN* in human colonic SMCs significantly attenuated contractile gene expression including *MYLK* and decreased SMC contractility. Luciferase reporter assays showed that *CARMN* enhances the transactivation activity of the master regulator of SMC contractile phenotype, myocardin, thereby maintaining the GI SMC myogenic program.

**Conclusion:** Our data suggest that *Carmn* is indispensable for maintaining GI SMC contractile function in mice, and that loss of function of *CARMN* may contribute to human visceral myopathy. To our knowledge this is the first study showing an essential role of lncRNA in the regulation of visceral SMC phenotype.

## Introduction

The major functions of the gastrointestinal (GI) tract are the digestion of food, absorption of nutrients and removal of waste. To accomplish these critical functions, smooth muscle cells (SMCs) populating the walls of the GI tracts generate the peristaltic forces that efficiently move food from one segment to the next or to retain food in one region until digestion and absorption are completed ^**1**^. Loss of GI motility is a characteristic of several smooth muscle-motility diseases including chronic intestinal pseudo-obstruction (CIPO) which is characterized by impaired contractility and functional intestinal obstruction ^**2**, **3**^. Smooth muscle-motility disorders have been shown to be caused by genetic mutations resulting in loss of function of SMC-contractile genes such as *MYH11* ^**4**^, *MYLK* ^**5**^, *LMOD1* ^**6**^, *ACTG2* ^**7**^, *ACTA2* ^**8**^ and *MYL9* ^**9**^ or the inactivation of their upstream regulatory transcription factors, such as *Srf* ^**10**, **11**^ and *Myocd* ^**12**^, two master regulators that form a complex binding to CArG elements within SMC-contractile gene loci ^**13**^. The etiology of CIPO is multifactorial, and the importance of epigenetic regulators such as long non-coding RNAs (lncRNAs) in this disease remains largely unexplored.

LncRNAs are a new class of RNA transcripts that have lengths exceeding 200 nucleotides but no apparent protein-coding potential. Accumulating evidence suggests that lncRNAs act as non-coding regulatory molecules that play critical roles in a variety of physiological and pathological conditions ^**14**, **15**^. Although several lncRNAs have been shown to contribute to SMC biology ^**16-23**^, these studies have exclusively focused on vascular SMCs. To our knowledge, thus far there has no reports on the functional role of lncRNAs in visceral SMCs.

In the current study, data mining of multiple independent publicly available omics datasets, revealed a novel lncRNA, *CARMN*, as the most abundantly expressed lncRNA in visceral SMCs of both human and mouse. *CARMN* was initially identified as an important regulator of cardiac differentiation in vitro ^24, 25^. Recently, we showed that *CARMN* is a highly abundant and conserved SMC-specific lncRNA, which plays a critical role in maintaining vascular SMC contractile phenotype ^26-28^. Unexpectedly, we found that both germline and SMC-specific inducible deletion of *Carmn* in mice resulted in premature lethality due to colonic pseudo-obstruction. These data suggest that *Carmn* is indispensable for GI function and to the best of our knowledge, it is the first lncRNA shown to have an important role in regulating GI motility. Our study further suggests that down-regulation or loss of function mutations of *CARMN* may contribute to CIPO in humans.

## Material and Methods

### Generation of *Carmn* global KO mice

The *Carmn* KO/GFP KI reporter mice were generated by insertion with a promoterless, reversed splicing acceptor/membrane-bound GFP gene trap cassette, which is flanked by 2 pairs of oppositely orientated Lox2272 sites and LoxP sites into intron 2 of the *Carmn* gene (referred to as conditional PFG allele) as we recently reported ^**26**^. Upon Cre-mediated deletion of exon 2 and inversion of the reversed GFP cassette (PFG to GFP), the inserted splicing acceptor will prematurely terminate *Carmn* expression by splicing exon 1 to the GFP cassette while turning on GFP expression under control of the endogenous *Carmn* promoter (referred to as GFP, or KO allele). GFP expression therefore faithfully displays endogenous *Carmn* expression *in vivo* while disrupting *Carmn* expression. To generate *Carmn* global KO (gKO) mice, we first crossed *Carmn*^PFG/PFG^ female mice with male mice ubiquitously expressing Cre (CMV-Cre, JAX, cat. #: 006054) ^29^ under the control of a human cytomegalovirus minimal promoter. The resultant heterozygous offspring mice were then intercrossed to obtain *Carmn*^GFP/GFP^ (gKO) mice, littermate WT (control) and *Carmn*^GFP/WT^ (Het) mice. Both male and female mice of the cohort of gKO and control mice are used.

### Generation of SMC-specific inducible *Carmn* KO mice

To generate SMC-specific *Carmn* inducible KO (iKO) mice, we first generated *Myh11*-CreER^T2+^; *Carmn*^PFG/WT^ mouse line, by crossing *Carmn*^PFG/PFG^ female mice with male mice expressing tamoxifen-inducible Cre driven by the SM-specific *Myh11* gene promoter (*Myh11*-CreER^T2+^) ^30^. Subsequently *Myh11*-CreER^T2+^; *Carmn*^PFG/WT^ male mice were bred with *Carmn*^PFG/PFG^ female mice to generate *Myh11*-CreER^T2+^; *Carmn*^PFG/PFG^ and *Myh11*-CreER^T2+^; *Carmn*^PFG/WT^ mice. The dual fluorescence reporter mTmG (membrane-targeted tandem dimer Tomato, mTomato or mT; membrane-targeted green fluorescent protein, mGFP or mG) mice (Stock No: 007676) ^31^ were purchased from the Jackson Laboratory. To generate the control mice, *Myh11*-CreER^T2+^ male mice were bred with homozygous mTmG^+/+^ female mice. The resulted *Myh11*-CreER^T2+^; mTmG^+/-^ SMC-lineage tracing mice were used as control to exclude potential cytotoxicity caused by ectopic expression of GFP and Cre in SMCs of *Carmn* iKO mice. To delete *Carmn* specifically in SMCs, 8-10 weeks old male *Myh11*-CreER^T2+^; mTmG^+/-^ (Control), *Myh11*-CreER^T2+^; *Carmn*^PFG/PFG^ (iKO) and *Myh11*-CreER^T2+^; *Carmn*^PFG/WT^ (iHet) mice were intraperitoneally injected with tamoxifen (1mg/mouse/IP) for 2 rounds of 5 days each, with a 2 days’ break in between. Only male mice are used in this study because *Myh11*-CreER^T2+^ transgene is only located in the Y chromosome ^30^. All mice used in this study are maintained on a C57BL/6J background. The primers for mouse genotyping are listed in **Supplemental Table 1**. The use of experimental animals has been approved by the Institutional Animal Care and Use Committee and Biosafety committee at Augusta University in accordance with NIH guidelines.

### Statistical Analysis

GraphPad Prism (version 9.2.0) was used for the statistical analysis. All data are expressed as mean ± SEM of at least 3 independent experiments. Tests used for statistical significance evaluations are specified in figure legends. An unpaired 2-tailed *t* test was used for data involving 2 groups only. Two-way analysis of variance was used for data involving >2 groups. Values of p < 0.05 were considered statistically significant, except for analysis of omics data, which used false discovery rate (FDR)-adjusted p < 0.05 as the threshold for statistical significance.

**A Detailed description of methods is provided in the Online Supplemental Material.**

## Results

### Identification of *CARMN* as a SMC-specific lncRNA in the human and mouse gut

To begin to screen for potentially important lncRNAs in GI tract, we first identified the most abundant lncRNAs that are expressed in human colon and small intestine tissues by analyzing the publicly available GTEx database ^**32**^. This analysis identified the top 10 highly enriched lncRNAs in adult human gut tissues (**Figure 1A**). To further examine the expression of these lncRNAs at a single cell level, we re-analyzed the single-cell transcriptome dataset of 62,849 cells isolated from duodenum, jejunum, ileum and colon of 6-10 weeks’ post-conception embryonic human gut (https://www.gutcellatlas.org) ^**33**^. Based on the gene expression signature of each cell, cells were divided into 45 clusters from 24 cell types (**Figure 1B** and **Supplemental Figure 1A**). UMAP (Uniform Manifold Approximation and Projection) visualization of gene expression revealed that among the top 10 most abundant lncRNAs, *CARMN* is the most restrictedly expressed lncRNA in SMCs, whose expression pattern is almost identical to that of the canonical SMC markers, such as *MYH11, MYLK, ACTA2, LMOD1, TAGLN* and *CNN1* and the SMC-specific transcription cofactor *MYOCD* ^**27**, **28**, **34**^ (**Figure 1C-D** and **Supplemental Figure1B**). In addition to SMCs, *CARMN* is also expressed in myofibroblasts and pericytes, both of which exhibit some extent of SMC features (**Figure 1D**). To examine the expression of *CARMN* in adult human colon tissues, we next de novo analyzed the scRNA-seq dataset that was generated from 16 adult human colons (GSE156905) ^**35**^. UMAP visualization revealed that, the expression of *CARMN* is specific in SMCs, paralleling with the expression of well-known SM-specific markers *MYH11* and *MYLK* (**Figure 1E** and **Supplemental Figure 1C**). Although bulk RNA-seq showed the lncRNAs *MALAT1, NEAT1* and *PGM5-AS1* are highly expressed in adult human gut (**Figure 1A**), none of them display a SMC-specific expression pattern as *CARMN* as revealed by the scRNA-seq analysis (**Figure 1C & E**). Furthermore, re-analyzing scRNA-seq dataset of adult mouse ileum and colon revealed that *Carmn* is restricted to SMCs, similar to the expression pattern of the well-known SMC-markers *Myh11* and *Mylk* and the SMC-specific transcription co-factor *Myocd* (**Supplemental Figure 1D**).

**Figure 1.**
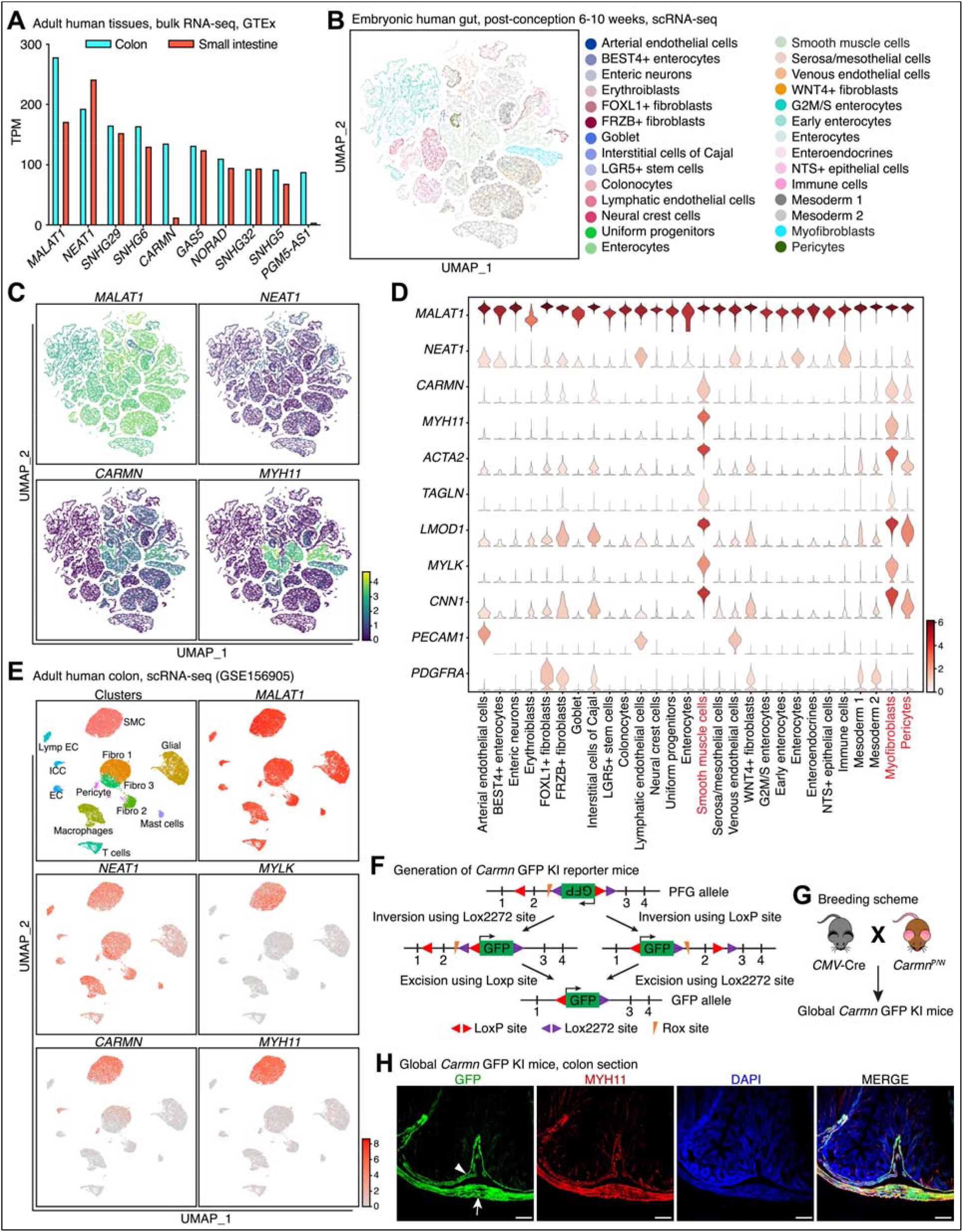
Identification of *CARMN* as a SMC-specific lncRNA in human and mouse GI tissues. (**A**) Analysis of the GTEx database showing the top 10 most abundant lncRNAs in human colonic and intestinal tissues. TPM: Transcripts per million. (**B**) UMAP (uniform manifold approximation and projection) plot to visualize the cell types revealed by the scRNA-seq analysis on developing human gut tissues (6-10 weeks post-conception) including duodenum, jejunum, ileum and colon. (**C**) UMAP visualization of gene expression for *MALTA1, NEAT1, CARMN* and *MYH11* in embryonic human gut revealed by the scRNA-seq analysis. The color scale on the bottom right corner indicates the expression levels. (**D**) Violin plot for the expression of selected genes (*MALAT1, NEAT1, CARMN, MYH11, ACTA2, TAGLN, LMOD1, MYLK, CNN1, PECAM1* and *PDGFRA*) in different cell types identified in embryonic human gut. *CARMN* is shown to be specifically expressed in SMCs, myofibroblasts and pericytes (in red). (**E**) UMAP plot to visualize cell types and the expression of *MALAT1, NEAT1, CARMN, MYLK* and *MYH11* in adult human colon tissues as revealed by the scRNA-seq analysis. (**F**) Strategy used to generate the *Carmn* GFP KI mouse model. (**G**) The breeding strategy to generate global *Carmn* GFP KI reporter mice by crossing the conditional *Carmn* heterozygous mice (P/W) with the global Cre mouse line driven by CMV promoter. (**H**) Direct visualization of GFP and immunostaining of MYH11 to examine the specific cellular localization of GFP in colon tissue dissected from K/W (Het) mice. Arrow and arrowhead denote muscularis externa SMCs and mucosa muscularis SMCs, respectively. Scale bar: 20 µm.

To validate and visualize the expression of *Carmn* in mice, we generated a *Carmn* KO/GFP KI mouse model with insertion of a promoterless, reversed splicing acceptor/membrane-bound GFP gene trap cassette, which is flanked by two pairs of oppositely orientated *Lox2272* sites and *LoxP* sites into the intron between exon 2 and 3 in mouse *Carmn* gene locus (PFG allele) ^26^ (**Figure 1F**). Cre-mediated recombination leads to inversion of the reversed GFP cassette (PFG to GFP, KO allele) and deletion of *Carmn* exon 2. Hence, *Carmn* transcription will be prematurely terminated by splicing of exon 1 to the splice acceptor of the GFP cassette, resulting in GFP expression driven by the endogenous *Carmn* gene promoter. To trace *Carmn* expression *in vivo*, we crossed the female *Carmn*^PFG/WT^ (*Carmn*^P/W^) mice with male mice expressing global Cre to invert the GFP cassette (**Figure 1G**). Although ubiquitous Cre recombines the PFG allele globally, data from the direct visualization of GFP fluorescence and the immune-fluorescence staining of the SM-marker MYH11 in K/W mouse colon tissue revealed that *Carmn* specifically expresses in MYH11 positive SMCs (**Figure 1H**). Similar findings were also observed in the section of jejunum from the K/W mouse (**Supplemental Figure 1E**). Taken together, data from these unbiased in silico analysis and from the novel *Carmn* KI GFP reporter mouse model demonstrate that *CARMN* is a lncRNA abundantly and specifically expressed in visceral SMCs in both human and mouse gut.

### Global deletion of *Carmn* in mice results in premature death due to the lethal colonic pseudo-obstruction

We next sought to explore functional role of *Carmn in vivo* by generating *Carmn* global KO (gKO) mice via intercrossing the female K/W and male K/W mice. The obtained WT and heterozygotes (Het) littermates serve as controls (**Figure 2A**). During the course of intercrossing *Carmn* Het mice, we unexpectedly observed approximate 60% of pregnant dams develop dystocia phenotype, suggesting haploinsufficiency of *Carmn* causes an impotency of uterine contraction to ensure successful parturition (**Figure 2B-C**). *Carmn* gKO mice were born at expected Mendelian ratio without gross abnormalities. However, on the weaning day of postnatal day 21, the gKO mice showed unhealthy appearance and smaller body size (**Figure 2D**) with significantly lower body weight compared to control mice (**Figure 2E**). Unexpectedly, after weaning and switching to the normal chow diet, approximate 60% of gKO mice exhibited an acute lethality within 1 week with all remaining gKO mice dying within 15 months. The gKO mice displayed an enlarged abdomen caused by dilated duodenum, jejunum and colon tracts due to infilled air and accumulation of stool (**Figure 2G**). Photograph of the isolated GI tracts further revealed that caecum and colon segments are the major distended parts with accumulation of feces in gKO mice (**Figure 2H**), suggesting a lethal colonic pseudo-obstruction. No apparent phenotype was observed in bladder, despite the high expression of *Carmn* in this tissue (**Figure 2G**). Histological analysis by HE staining further revealed severe dilation of colon and thinning muscle layers of gKO mice compared to control mice (**Figure 2I-J**). Similar results were also observed in the jejunum (**Supplemental Figure 2A-B**). Moreover, ultrastructural examination by transmission electron microscopy revealed that *Carmn* KO SMCs in both colon and jejunum display markedly abnormal structures, including damaged endoplasmic reticulum with dramatically dilated lumen and massive membrane laminated vacuoles with embedded membranes and organelles such as mitochondria, suggesting a stressed and degenerative phenotype of SMCs (**Figure 2I** and **Supplemental Figure 2C**). Taken together, these data demonstrate that global deletion of *Carmn* causes lethal colonic pseudo-obstruction and dystocia, and suggest that *Carmn* is indispensable for the GI and uterine function.

**Figure 2.**
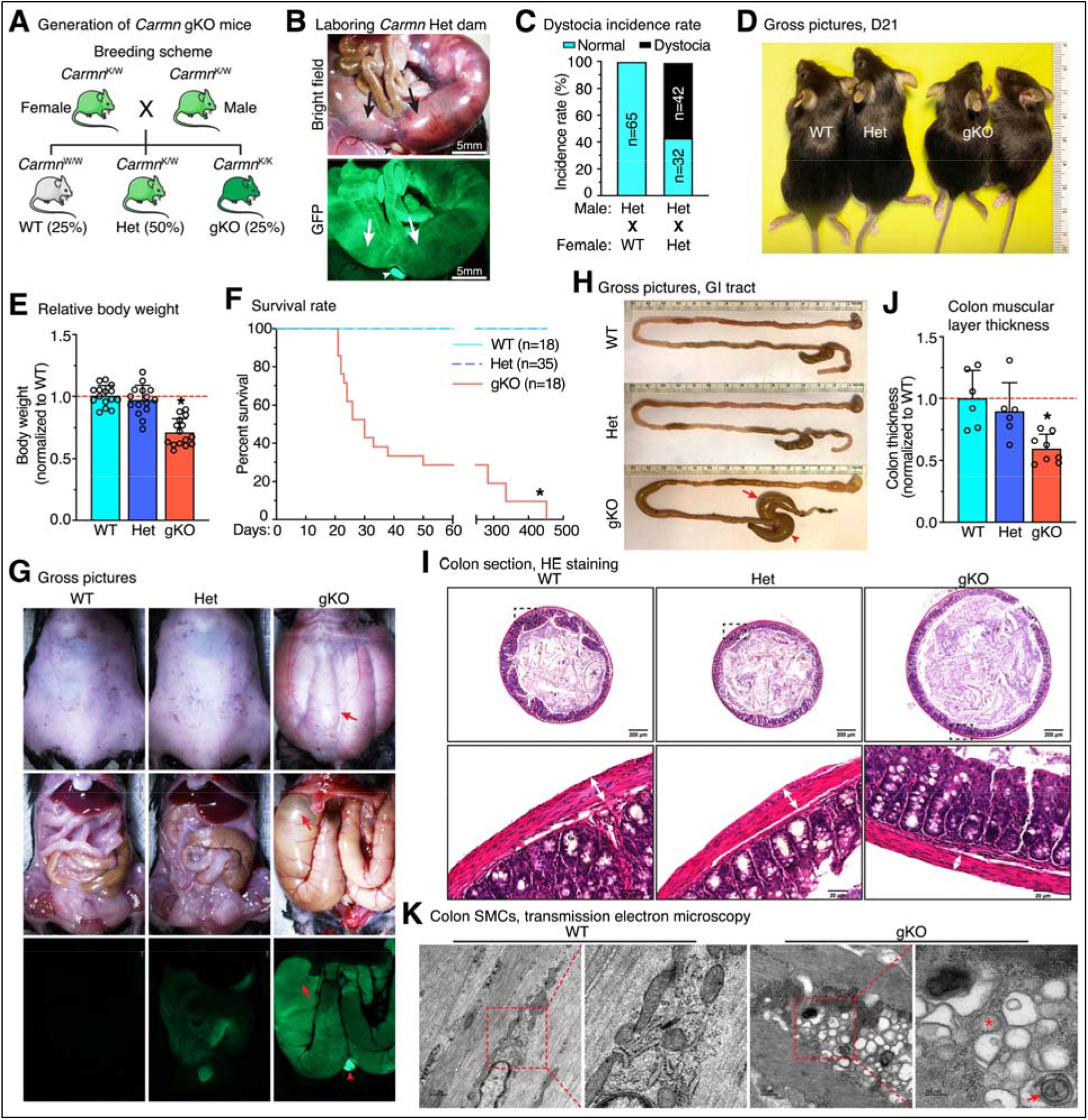
Global deletion of *Carmn* in mice causes lethal colonic pseudo-obstruction. (**A**) The breeding strategy to generate global *Carmn* KO mice by intercrossing *Carmn* Het (K/W) mice. (**B**) The representative picture of uterus from a laboring *Carmn* Het female mouse under bright field (upper panel) or GFP channel (bottom panel) to show the dystocia phenotype. Arrows and arrowhead denote the trapped embryos in the uterine tract and bladder, respectively. (**C**) The dystocia incident rate of the laboring WT and *Carmn* Het female mice crossed with *Carmn* Het male mice. (**D**) Gross pictures of littermates of WT, Het and *Carmn* gKO mice to show the smaller body size of the gKO mice on postnatal day (D) 21. (**E**) The relative body weight of the littermates WT, Het and *Carmn* gKO mice on postnatal D21. N = 15; *p < 0.05; Unpaired Student *t* test. (**F**) Kaplan-Meier survival analysis of WT, Het and *Carmn* gKO mice. N = 18; *p < 0.05; Log-rank (Mantel-Cox) test. (**G**) *Carmn* gKO mice exhibit abdominal distension (top panel) and dramatically enlarged intestine with accumulated air (arrow) compared to WT and Het littermates. The pictures were acquired under a dissecting scope with bright field (top and middle panels, before and after opening abdominal cavity, respectively) and GFP channel (bottom panel after opening the abdominal cavity). The arrowhead points to the bladder. (**H**) Gross pictures of the GI tract isolated from WT, Het and *Carmn* gKO mice. The dramatically affected GI segments in the *Carmn* gKO mice are the cecum (arrowhead) and colon (arrow). (**I**) Hematoxylin and eosin (HE) staining on the transverse sections of colon from WT, Het and *Carmn* gKO mice. The boxed area is magnified on the bottom of each respective panel. (**J**) The relative thickness of colon muscularis layers (double-head arrows shown in “**I**”) was measured and plotted. N = 6-8; *p < 0.05; Unpaired Student *t* test. (**K**) Representative transmission electron microscopy (TEM) images of control and *Carmn* gKO colon show the enlarged endoplasmic reticulum lumen (asterisk), autophagic vesicles with lamination (arrow) in gKO mouse colon, indicating that the *Carmn* deficient SMCs are under stress and are degenerative. The boxed area is magnified on the right.

### Inducible SMC-specific deletion of *Carmn* in adult mice recapitulates the lethal phenotype of *Carmn* gKO mice

To uncover the functional role of *Carmn* specifically in adult SMCs, we generated *Carmn* inducible SMC-specific KO mice by crossing tamoxifen (TAM)-inducible transgenic mice carrying SMC-specific gene *Myh11* promoter driven CreER^T2^ (*Myh11*-CreER^T2^) with *Carmn*^PFG/PFG^ mice (referred to as iKO mice hereafter, **Figure 3A**). To exclude potential cytotoxicity caused by ectopic expression of GFP and Cre in SMCs of SM-specific *Carmn* iKO mice, we used the SM-lineage tracing mice (referred to as WT control mice hereafter) as control in which Cre and membrane-tagged GFP specifically express in SMCs as iKO mice by crossing *Myh11*-CreER^T2^ mice with Rosa26-mTmG dual fluorescence reporter mice (**Figure 3B**). We also included a cohort of mice in which a single *Carmn* allele is specifically deleted in adult SMCs as an additional control (referred to as iHet hereafter). TAM was peritoneally injected into these 3 cohorts of 8-10 weeks old male mice for 10 times and then the mouse body weight was monitored (**Figure 3C**). We found at day 62 post the first TAM injection, the body weight of iKO mice is significantly reduced compared to the WT and iHet mice. We observed that iKO mice start to display mortality at day 33, and none of them can survival beyond 130 days after the first TAM injection (**Figure 3D**). Dissecting the iKO mice at day 30 post the 1^st^ TAM injection revealed almost identical the pathological events observed in gKO mice, including drastic distention of GI tract that is infilled with abundant air and feces, especially in the cecum and colon segments (**Figure 3E-F**). Histological analysis of colon and jejunum of iKO mice showed dilation and thinner muscular layer compared to the WT and iHet control mice, similar findings as gKO mice (**Figure 3G-H** and **Supplemental Figure 3A-B**). Ultrastructural images revealed a similar severe degenerative phenotype of SMCs in both colon and jejunum of iKO mice as observed in gKO mice (**Figure 3I** and **Supplemental Figure 3C**). Taken together, these data demonstrate that deletion of *Carmn* specifically in postnatal SMCs of adult mice recapitulates the lethal colonic pseudo-obstruction phenotype occurred in *Carmn* gKO mice, leading to premature lethality.

**Figure 3.**
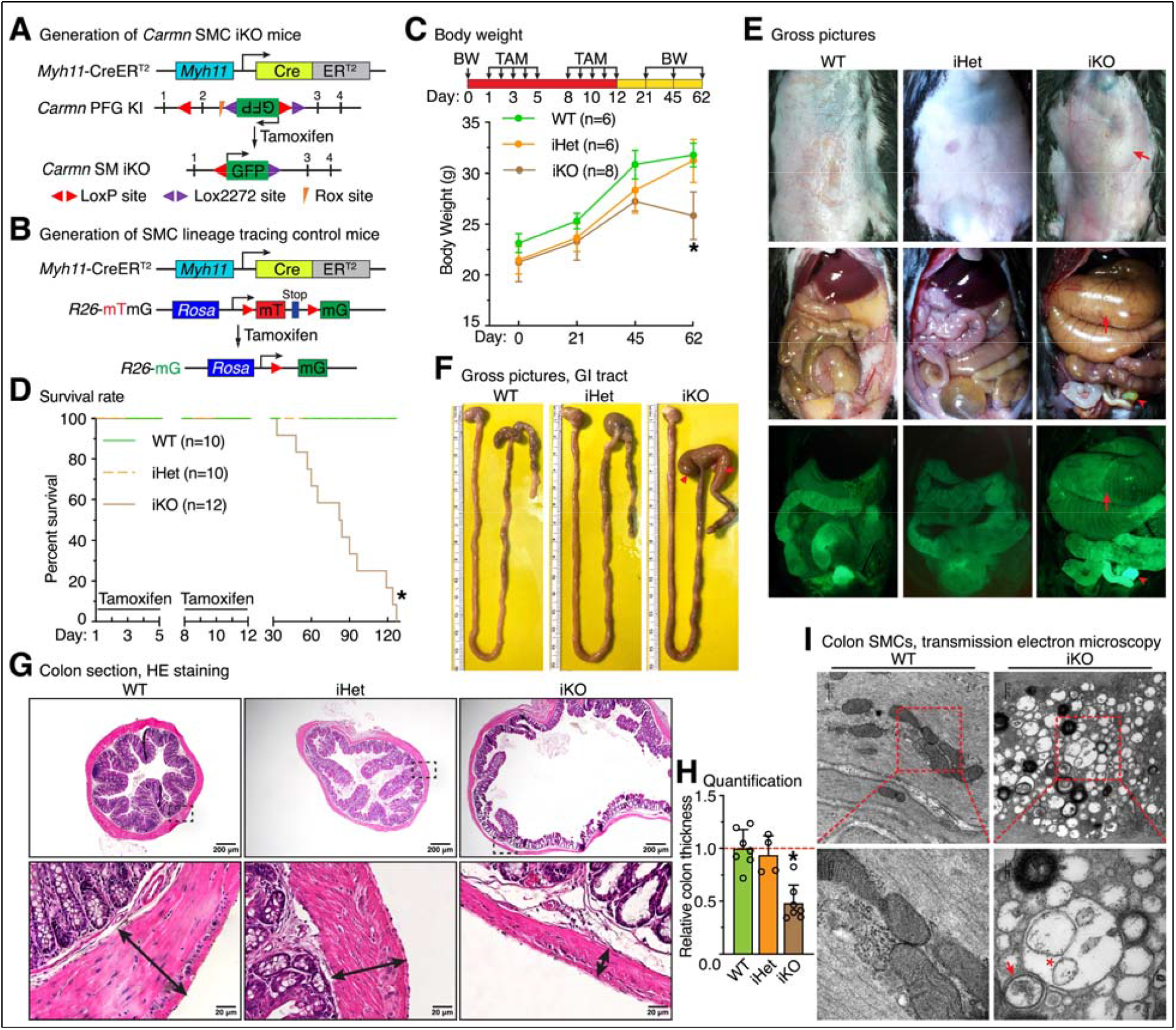
SMC-specific deletion of *Carmn* in adult mice causes lethal colonic pseudo-obstruction, phenocopying that of *Carmn* gKO mice. (**A-B**) Schematic diagram showing SMC-specific gene *Myh11* promoter-driven CreER^T2^ mediated inducible SMC-specific deletion of *Carmn* in adult mice (iKO) by injection of tamoxifen into adult male *Myh11*-CreER^T2+^; *Carmn*^F/F^ mice and **(B)** lineage tracing control mice (WT) by injection of tamoxifen into adult male *Myh11*-CreER^T2+^; *R26*-mTmG^+/-^ mice, respectively. The mTmG reporter mice contain a dual fluorescence reporter transgene integrated into the *Rosa26* locus. Tamoxifen (TAM)-dependent Cre activation will remove membrane localized Tomato (mT)-stop cassette thereby inactivating mT expression, while membrane localized GFP (mG) will be switched on to permanently mark the Cre-expressing cells and their progeny. (**C**) The body weight of the WT, iHet and iKO mice were measured before and after TAM injection at the time indicated. N = 6-8; *p < 0.05; 2-way analysis of variance followed by post hoc testing within day. (**D**) Kaplan-Meier survival analysis of WT, iHet and iKO mice after two rounds of intraperitoneal injections with TAM. N = 10-12; *p < 0.05; Log-rank (Mantel-Cox) test. (**E**) *Carmn* iKO mice exhibit abdominal distension and dramatic intestinal enlargement (arrows) compared to WT and iHet mice. Arrowhead denotes the bladder. The pictures were acquired under a dissecting scope with bright field (top and middle panels, before and after opening abdominal cavity, respectively) and GFP channel (bottom panel after opening the abdominal cavity). (**F**) Pictures of the GI tract isolated from WT, iHet and iKO mice on day 30 post the 1^st^ TAM injection. The cecum (arrowhead) and colon (arrow) are the most affected parts in the GI tract of *Carmn* iKO mice. (**G**) HE staining on the transverse sections of colon from WT, iHet and iKO mice. The boxed area is magnified on the bottom. (**H**) The thicknesses of muscular layers of colon were measured (double-head arrows in “**G**”) and plotted. N = 4-7; *p < 0.05; Unpaired Student *t* test. (**I**) Representative transmission electron microscopy images demonstrate formation of a significant amount of autophagic vesicles (arrow) and a large vesicle containing several small vesicles (asterisk) in *Carmn* iKO colon SMCs. The boxed area is magnified on the bottom.

### *Carmn* deficiency impairs GI motility and colonic contractility in mice

Previous studies have shown that the impairment of visceral SMC contraction in the GI tract is responsible for colonic pseudo-obstruction associated with CIPO in humans ^**5**, **6**, **8**, **9**^. To directly assess the contractile function of GI tract of gKO and iKO mice *in vivo*, the whole-gut transit time from food intake to excretion were measured as a functional readout of GI motility. We found that the whole-gut transit time was significantly increased in both *Carmn* KO mouse models (gKO and iKO) compared to their respective control mice (**Figure 4A-B**), indicating that *Carmn* is indispensable for the GI motility. The impairment of gut motility in iKO mice resulted in larger stool size in diameter and darker color stool (**Figure 4C-D**). As the phenotype mainly developed in the colon segment, next we measured the colonic contractile activity ex vivo using myography. We found that the spontaneous colonic contractile activity in both *Carmn* gKO and iKO mice is significantly attenuated compared to that of control mice (**Figure 4E-F** and **Supplemental 4A-B**). Moreover, the amplitude of colonic contraction elicited by stimulation of KCl (60 mM) or the muscarinic agonist Carbachol (1 µM Cch), both of which can initiate smooth muscle contraction through MYLK-dependent phosphorylation of myosin regulatory light chain (RLC) ^**36**, **37**^, was significantly impaired in *Carmn* gKO and iKO mice compared to their respective control mice (**Figure 4G-N** and **Supplemental Figure 4C-F**). In summary, these data indicate that *Carmn* is indispensable for GI motility and colonic contractility.

**Figure 4.**
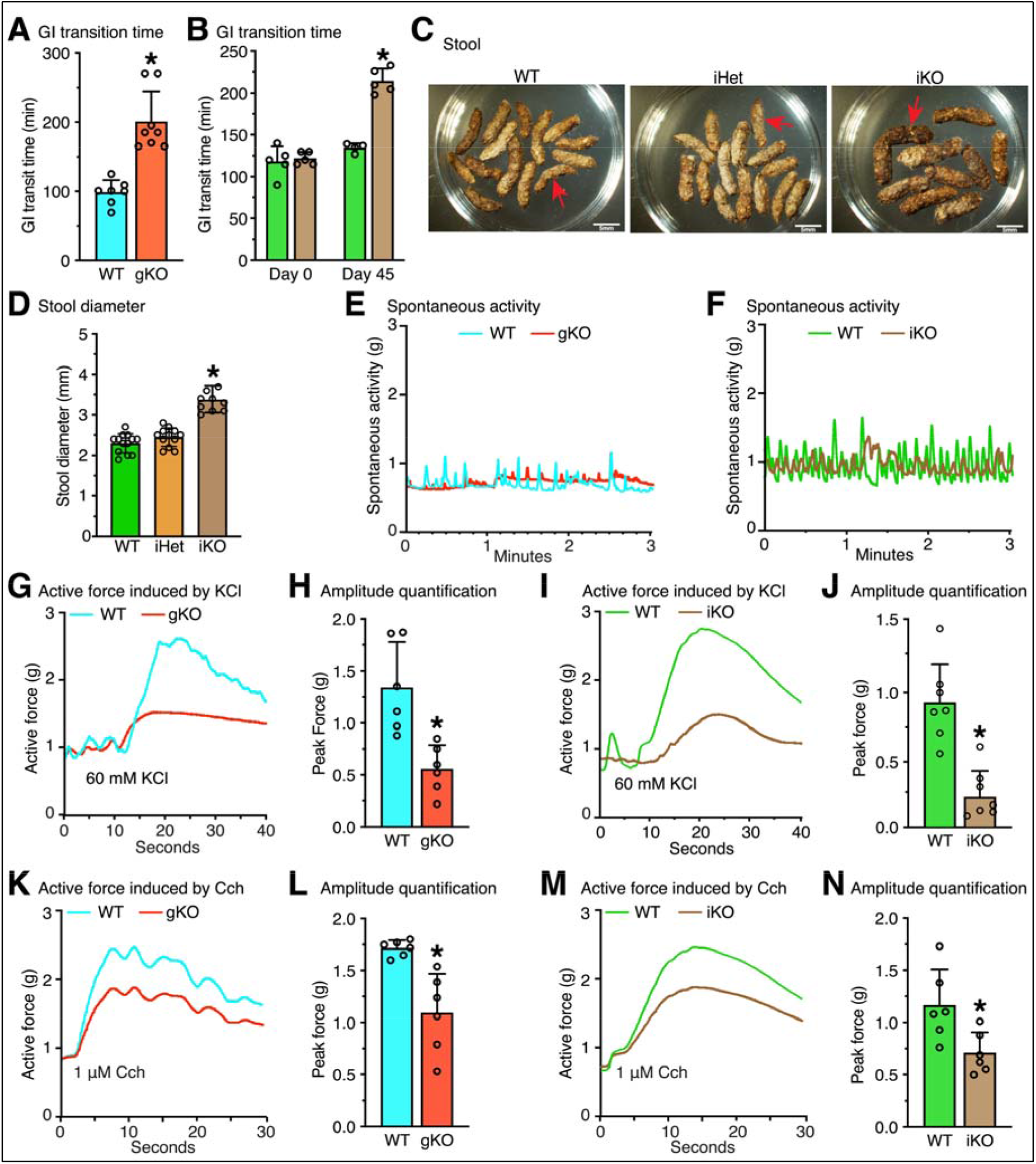
*Carmn* deficiency impairs GI motility and colonic contractility. (**A**) Whole-gut transit time for WT and *Carmn* gKO mice. N = 7-8; *p < 0.05; Unpaired Student *t* test. (**B**) Whole-gut transit time for WT and *Carmn* iKO mice at day 0 before and at day 45 after TAM injection. N = 5; *p < 0.05; 2-way analysis of variance followed by post hoc testing within day. (**C-D**) Stools were collected from WT control, *Carmn* iHet and iKO mice on day 30 post the 1^st^ TAM injection. The stool diameter was then measured and plotted. N = 9-13; *p < 0.05; Unpaired Student t test. (**E-F**) Representative spontaneous contraction recordings in the colonic rings from *Carmn* gKO (**E**) or (**F**) iKO mice their respective WT control mice. (**G-J**) Representative recordings of contraction induced by 60 mM KCl on colonic rings from (**G**) *Carmn* gKO or (**I**) iKO mice and their respective WT control mice. Quantitative analysis of peak force induced by KCl treatment on *Carmn* gKO or iKO mouse colonic rings are shown in “**H**” or “**J**”, respectively. N = 6-7; *p < 0.05; Unpaired Student *t* test. (**K-N**) Representative recordings of contraction induced by 1 µM Cch (Carbachol, a muscarinic receptor agonist) on colonic rings from (**K**) *Carmn* gKO or (**M**) iKO mice and their respective WT control mice. The peak force induced by Cch treatment on *Carmn* gKO or iKO mouse colonic rings are shown in “**L**” or “**N**”, respectively. N = 6; *p < 0.05; Unpaired Student *t* test.

### *Carmn* deficiency induces visceral SMC de-differentiation

To understand the molecular mechanism that underlies *Carmn* deficiency-induced colonic pseudo-obstruction phenotype, bulk RNA-sequencing (RNA-seq) was performed on the muscularis isolated from colon and jejunum tissues of *Carmn* iKO and control mice (**Figure 5A**). Data from the RNA-seq analysis demonstrated that *Carmn* iKO resulted in 318 significantly down-regulated genes and 243 up-regulated genes in colon muscularis (**Figure 5B** and **Supplemental Table 2**), while 561 significantly down-regulated genes and 635 up-regulated genes in jejunum muscularis, respectively (**Figure 5C** and **Supplemental Table 2**). Gene ontology (GO) analysis of down-regulated genes in colon muscularis revealed that *Carmn* deficiency significantly attenuates the genes involved in maintaining the muscle homeostasis, such as muscle contraction, muscle tissue development and muscle cell differentiation (**Figure 5D & F**). In contrast, up-regulated genes in colon muscularis are significantly over-represented in pathways responding to tissue repair such as matrix remodeling, wound healing, and inflammatory process including response to lipopolysaccharide (**Figure 5E-F**). In *Carmn* KO jejunum muscularis, the down-regulated genes are involved in metabolic processes (**Supplemental 5A**), while the up-regulated genes are associated with immune cell infiltration and inflammatory reaction (**Supplemental 5B**), a similar finding observed in iKO colon tissues. To better understand the potential genes directly regulated by *Carmn*, we further compared the overlapping differentially expressed genes in both colon and jejunum muscularis. The results showed by the Venn diagram demonstrated that there are 136 overlapping downregulated and 128 overlapping upregulated genes between *Carmn* iKO colon and jejunum muscularis, respectively (**Figure 5G**). Strikingly, GO analysis of the overlapping down-regulated genes revealed that these over-represented genes belonging to muscle contraction and development, almost identical to the results observed in colon. These data collectively suggest that the down-regulation of these contractile genes is a primary cause for the hypomotility of GI tracts observed in *Carmn* deficient mice (**Figure 5H** and **Supplemental Figure C**). Consistently, GO analysis of the up-regulated genes in common revealed the shared pathways are involved in inflammatory response and matrix remodeling (**Supplemental Figure 5D-E**), implicating a secondary effect to ileus paralytics induced by deficiency of *Carmn*. To further pinpoint the key targets of *Camrn* with critical roles in maintaining colonic SMC homeostasis, we analyzed the common genes involved in multiple smooth muscle homeostasis signaling pathways. This analysis revealed that *Mylk* is the only overlapping gene, which has been shown to be the principal regulator of the myosin II molecular motor in the initiation of smooth muscle contraction essential for normal gastrointestinal motility (**Figure 5I**) ^**38**^. Taken together, the integrative analysis of the bulk RNA-seq datasets on both colon and jejunum muscular layers suggest that *Carmn* deficiency disrupts the homeostatic expression of muscle contractile genes such as *Mylk* while promoting expression of genes involved in response to tissue injury and inflammation.

**Figure 5.**
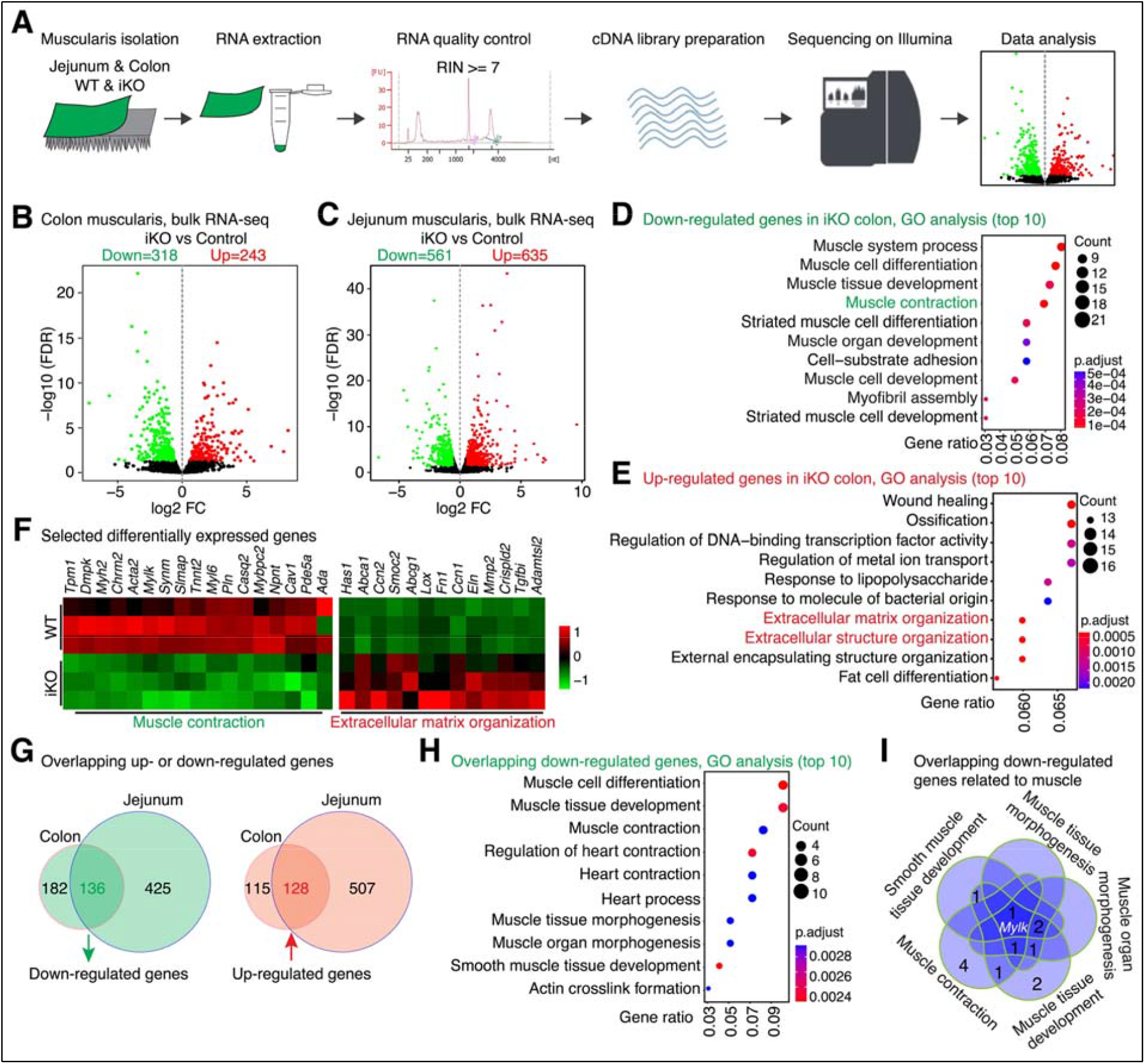
*Carmn* deficiency down-regulates the expression of genes involved in muscle contraction while increasing the expression of genes related to extracellular matrix remodeling. (**A**) The workflow of bulk RNA-seq on colon and jejunum muscularis isolated from WT and iKO *Carmn* KO mice. (**B**) Volcano plots depicting the significantly down-regulated and up-regulated genes between *Carmn* iKO and WT mouse colon and (**C**) jejunum muscularis. FDR: False Discovery Rate. FC: Fold Change. (**D**) GO analysis showing that the 318 down-regulated genes in *Carmn* iKO colon muscularis are significantly enriched in the biological processes related to muscle contraction and differentiation (**E**) while the significantly up-regulated genes are enriched in the functional categories involved in the wound healing and extracellular matrix remodeling. The top 10 most significantly enriched GO biological processes are shown. (**F**) Heat map showing the differential expression of selected genes involved in muscle contraction and extracellular matrix organization. (**G**) Venn diagrams showing the overlapping down-regulated (left panel) or up-regulated (right panel) genes between colon and jejunum muscularis in *Carmn* iKO mice. (**H**) GO analysis showing that the 136 overlapping down-regulated genes in both colon and jejunum muscularis of *Carmn* iKO mice are significantly enriched in the biological processes related to muscle contraction. The top 10 most significantly GO biological processes were shown. (**I**) Venn diagram showing the identification of overlapping down-regulated genes involved in biological processes of muscle function.

### *Carmn* deficiency disrupts homeostatic SMC contractile gene program and compromises the cell-cell communication in the colonic muscularis

To obtain a comprehensive understanding of changes in transcriptome of *Carmn* deficient SMCs at the single cell level, we isolated cell nuclei from colonic muscularis of *Carmn* iKO and control mice and performed snRNA-seq (**Supplemental Figure 6A**). UMAP analysis of 5,072 nuclei from both control and *Carmn* iKO samples revealed 17 distinct clusters with 13 cell types as defined by the cell type-specific gene expression profile (**Figure 6A** and **Supplemental Figure 6B-E**). As expected, *Carmn* was specifically detected in SMCs positive for the well-known SMC marker genes *Myh11* and *Mylk* and the SMC-specifically expressed transcription co-factor *Myocd* (**Figure 6A** and **Supplemental Figure 6E**). Cell composition analysis revealed that compared to control group, the percentage of SMCs, glial cells, Ctfr+ cells and Goblet cells is decreased, while the percentage of fibroblast, fibroblast-like, macrophage and T cells is increased in *Carmn* iKO samples (**Figure 6B**). The decreased SMCs, Ctfr+ cells and Goblet cells may contribute to the imbalance of colon muscularis homeostasis ^**39**^, while the increased fibroblast or fibroblast-like, macrophage and T cells may be responsible for the tissue repair and inflammatory response as secondary to the severe impairment of GI motility observed in *Carmn* iKO mice ^**40**, **41**^. To unravel the transcriptome changes of SMCs following *Carmn* deletion, we next conducted a differential analysis specifically in SMC clusters. This analysis identified 123 down-regulated and 265 up-regulated genes, respectively, in the SMCs of *Carmn* iKO compared to control (**Figure 6C** and **Supplemental Table 3**). GO analysis of the down-regulated genes in iKO SMCs revealed that *Carmn* deficiency significantly attenuates the genes involved in regulating muscle contraction and synapse organization (**Supplemental Figure 6F**), while the top 10 enriched pathways of the up-regulated genes are mainly responsible for extracellular matrix organization and adhesion (**Supplemental Figure 6G**), pathways tightly related with tissue repair ^**42**^. Consistent with the bulk RNA-seq data, genes related to muscle contraction including *Mylk* and *Chrm2* were among the significantly down-regulated genes while the matrix genes such as *Ccn2, Fn1* and *Eln* were significantly increased in the iKO SMCs, indicating a de-differentiation status of iKO SMCs (**Supplemental Figure 6H**). Analysis of the differentially expressed genes (DEGs) in colon muscularis revealed by bulk RNA-seq and in SMC clusters revealed by snRNA-seq uncovered 29 co-downregulated genes and 34 co-upregulated genes, respectively (**Figure 6D**). Intriguingly, GO analysis of these co-downregulated and co-upregulated genes showed that these genes are involved in the pathways regulating muscle contraction and extracellular matrix organization, respectively (**Figure 6E-F**). Taken together, results from the colon muscularis bulk RNA-seq and snRNA-seq unambiguously demonstrate that deletion of *Carmn* in SMCs leads to a de-differentiation phenotype of SMCs by down-regulating contractile genes while up-regulating genes involved in extracellular matrix remodeling.

**Figure 6.**
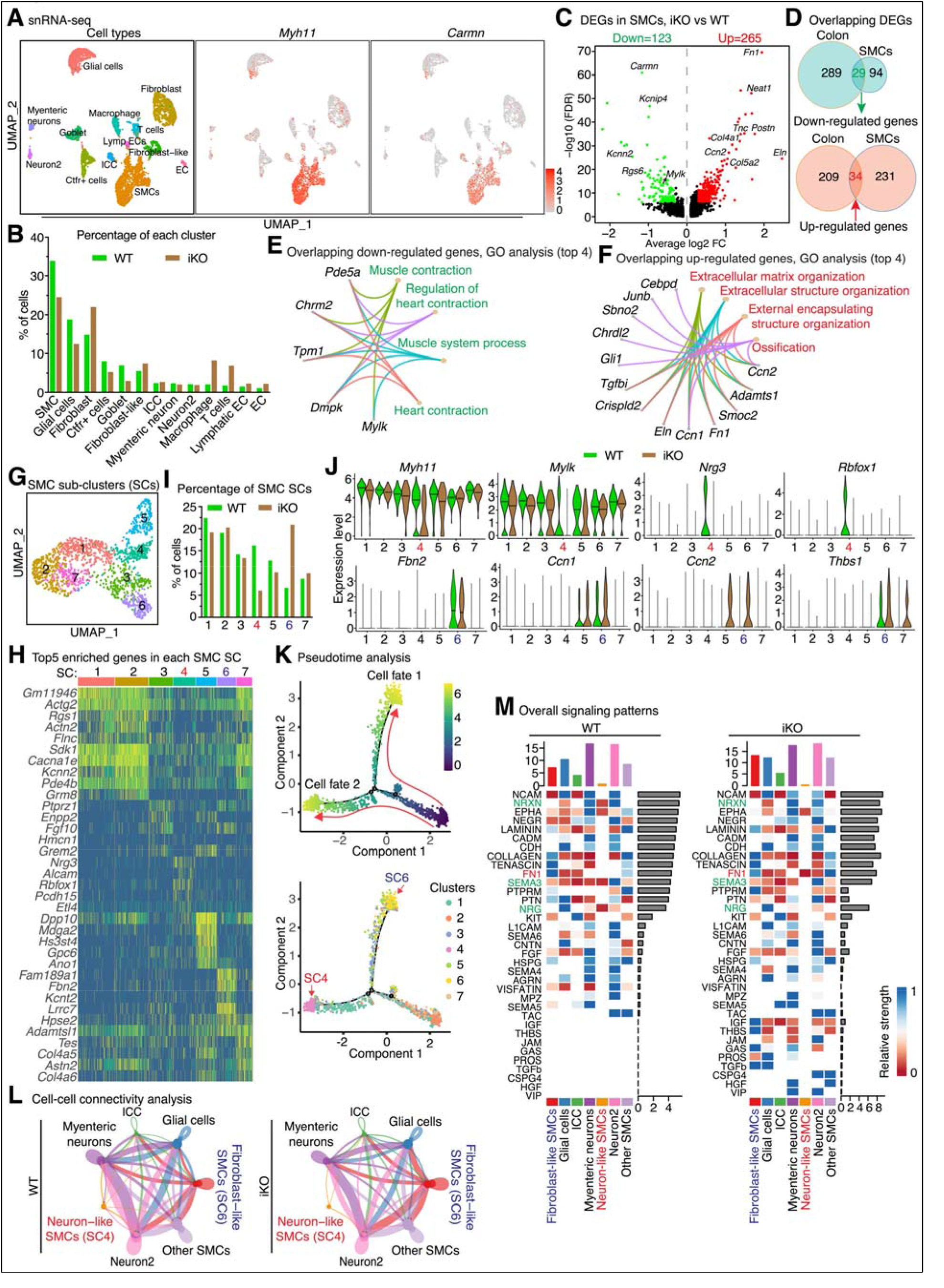
Single nucleus RNA-sequencing (snRNA-seq) identifies changes of SMC specific sub-clusters in colon muscularis of *Carmn* iKO mouse. (**A**) UMAP visualization of *Myh11* and *Carmn* expression in the cell types of colon muscularis revealed by snRNA-seq. (**B**) Percentage of each cell types in the colon muscularis from WT and *Carmn* iKO mice. (**C**) Volcano plot showing the differentially expressed genes (DEGs) specifically in SMCs between *Carmn* iKO and WT colon muscularis as revealed by the snRNA-seq. (**D**) Venn diagrams showing the overlapping down-regulated (upper panel) or up-regulated (bottom panel) genes in SMC clusters revealed by snRNA-seq and in colon muscularis identified by bulk RNA-seq in *Carmn* iKO mice. (**E**) GO analysis of the top 4 overlapping GO biological processes and involved genes for the down-(**F**) and up-regulated genes as shown in “**D**”. (**G**) UMAP plot showing 7 distinct SMC sub-clusters (SCs) in mouse colonic muscularis. (**H**) Heatmap showing the top 5 enriched genes in each SMC SC. (**I**) Percentage of each SMC SC in both *Carmn* iKO and WT mouse colonic SMCs. (**J**) Violin plots showing the expression level of a subset of genes in each SC of SMCs. (**K**) Pseudotime trajectory analysis showing the branching SMC fates (upper panel) and the fate transition of each SMC SC along the trajectory. (**L**) Cell-cell connectivity analysis showing the cell-cell interaction among all cell types identified in WT and *Carmn* iKO mouse colon muscularis. The line thickness and node size correspond to the number of cognate ligand-receptor expression and cell numbers, respectively. (**M**) Heatmaps showing the overall (both outgoing and incoming) signaling patterns of each cell type mediated by individual ligand signaling axes in both WT (left panel) and *Carmn* iKO (right panel) mouse colon muscularis. The NRXN, SEMA3 and NRG ligands-mediated signaling (in green) is lost while FN1 signal (in red) is gained in neuron-like SMCs (SC4) in *Carmn* iKO SMCs, compared to WT SMCs.

To define the sub-populations of SMCs in mouse colon in greater detail, we closely examined the transcriptome of the 1,529 SMCs identified in both iKO and control samples. Unsupervised Seurat-based clustering of those SMCs revealed 7 distinct sub-populations defined by cluster-specific marker genes (**Figure 6G-H**). Interestingly, we found that, compared to control WT SMCs, *Carmn* iKO SMCs contain a reduced percentage of sub-cluster 4 and a higher percentage of sub-cluster 6 (**Figure 6I**). Although all 7 SMC subpopulations are *Myh11* positive (the 1^st^ panel in the top row in **Figure 6J**), expression of *Myh11* gene is lower whereas expression *Mylk* gene (the 2^nd^ panel in the top row of **Figure 6J**) is nearly completely absent in the SMC sub-cluster 4 in *Carmn* iKO SMCs, compared to control. In contrast, the most significantly enriched 2 genes in the sub-cluster 4 SMCs but depleted in iKO SMCs are *Nrg3* (Neuregulin 3, a ligand binding to ERBB-family receptors) and *Rbfox1* (RNA binding protein, fox-1 homolog) (**Figure 6J**), both are neuron-enriched genes (**Supplemental Figure 6E)**. Notably, *NRG3* gene mutations in enteric neuron system cause Hirschsprung’s disease (HSCR) in humans ^**43**^, a birth defect that characterized with bowl hypoperistalsis. In contrast, SMCs in sub-cluster 6 are the only subset of SMCs positive of the fibroblast-specific gene *Fbn2* (the 1^st^ panel in the bottom row of **Figure 6J**). Furthermore, the most significantly upregulated genes in iKO SMC sub-cluster 6 are *Ccn1, Ccn2* and *Thbs1*, all of which are fibroblast-specific genes and are key mediators for smooth muscle extracellular matrix homeostasis (**Figure 6J**) ^**44**^. To better understand the transcriptome characteristics of the SMC sub-cluster 4 and 6, GO analysis of the enriched genes in both sub-clusters showed that the enriched genes are involved in the pathways regulating axonogenesis and matrix remodeling, respectively (**Supplemental Figure 7A-B**). Therefore, we refer to the sub-cluster 4 as neuron-like SMCs and sub-cluster 6 as fibroblast-like SMCs. To investigate the transition of SMC phenotypes, we applied the Monocle 2 algorithm to perform the single-cell pseudotime trajectory analysis ^**45-47**^. This analysis revealed SMCs form a continuous transition and progressively branched into two trajectories, ending with SMC sub-cluster 4 and 6, respectively (**Figure 6K** and **Supplemental Figure 7C**). Collectively, this comprehensive snRNA-seq analysis revealed that the SMCs in mouse colon muscularis can be divided into three groups: the normal contractile SMCs (clusters 1, 2, 3, 5, 7) on which *Carmn* deficiency has little effects, the novel neuron-like SMCs (sub-cluster 4) that is depleted in the *Carmn* iKO mice, and the novel fibroblast-like SMCs (sub-cluster 6) which is increased in the *Carmn* iKO SMCs. Thus, we hypothesize that the higher percentage of the fibroblast-like de-differentiated SMCs are responsible for tissue repair, while the reduced percentage of the neuron-like SMCs in *Carmn* iKO mice may impair the neurogenic motility that synergistically communicates with other cell types ^**48**, **49**^. To test this hypothesis, we performed cell-cell communication among these cells using CellChat ^**50**^. This analysis revealed that, compared with WT mice, neuron-like SMCs in *Carmn* iKO cells lost the cell-cell communication with neuron2, other contractile SMCs, glial cells, ICC and myenteric neurons (**Figure 6L** and **Supplemental Figure 7D**). The overall signaling patterns showed that compared to WT cells, *Carmn* iKO SMCs gain cellular signaling involved in SMC de-differentiated, such as IGF, THBS and TGFb, which closely correlated with the increased percentage of fibroblast-like SMCs in *Carmn* iKO mice (**Figure 6M**). In contrast, neuron-like SMCs lost the NRXN, SEMA3 and NRG signals while gaining FN1 ligand-mediated signal (**Figure 6M** and **Supplemental Figure 7E**). Taken together, the data from the snRNA-seq analysis demonstrate that *Carmn* deficiency in SMCs not only compromises the myogenic motility by reducing expression of contractile genes essential for SMC contraction but also impairs neurogenic motility by disrupting the cell-cell connectivity in the colonic muscularis, collectively contributing to the colonic pseudo-obstruction phenotype of *Carmn* iKO mice.

### Deletion of *Carmn* attenuates expression of genes essential for SMC contraction

Next we validated the bulk RNA-seq and snRNA-seq data by assessing the expression of contractile genes regulated by *Carmn*. Results from qRT-PCR and Western blotting assays demonstrated that MYLK is consistently down-regulated in both jejunum and colon muscularis of both gKO and iKO mice, compared to their respective control mice, suggesting that *Mylk* is a major target gene regulated by *Carmn* (**Figure 7A-F** and **Supplemental Figure 8A-B**). *Carmn* deficiency did not significantly affect the expression of the key transcription factor *Srf* and its cofactor *Myocd* (**Figure 7A** and **D**). IF staining further revealed that the MYLK immuno-fluorescence intensity in both colonic and intestinal muscular layers of both gKO and iKO mice are markedly weaker than that of controls (**Figure 7G-H** and **Supplemental Figure 7C-D**). Taken together, these data demonstrate that *Carmn* deficiency leads to decreased expression of the key SM-contractile gene *Mylk* that is essential for SM contraction, indicating that the lethal colonic pseudo-obstruction phenotype observed in *Carmn* KO mice is attributable to the decreased expression of *Mylk* gene in *Carmn* deficient SMCs.

**Figure 7.**
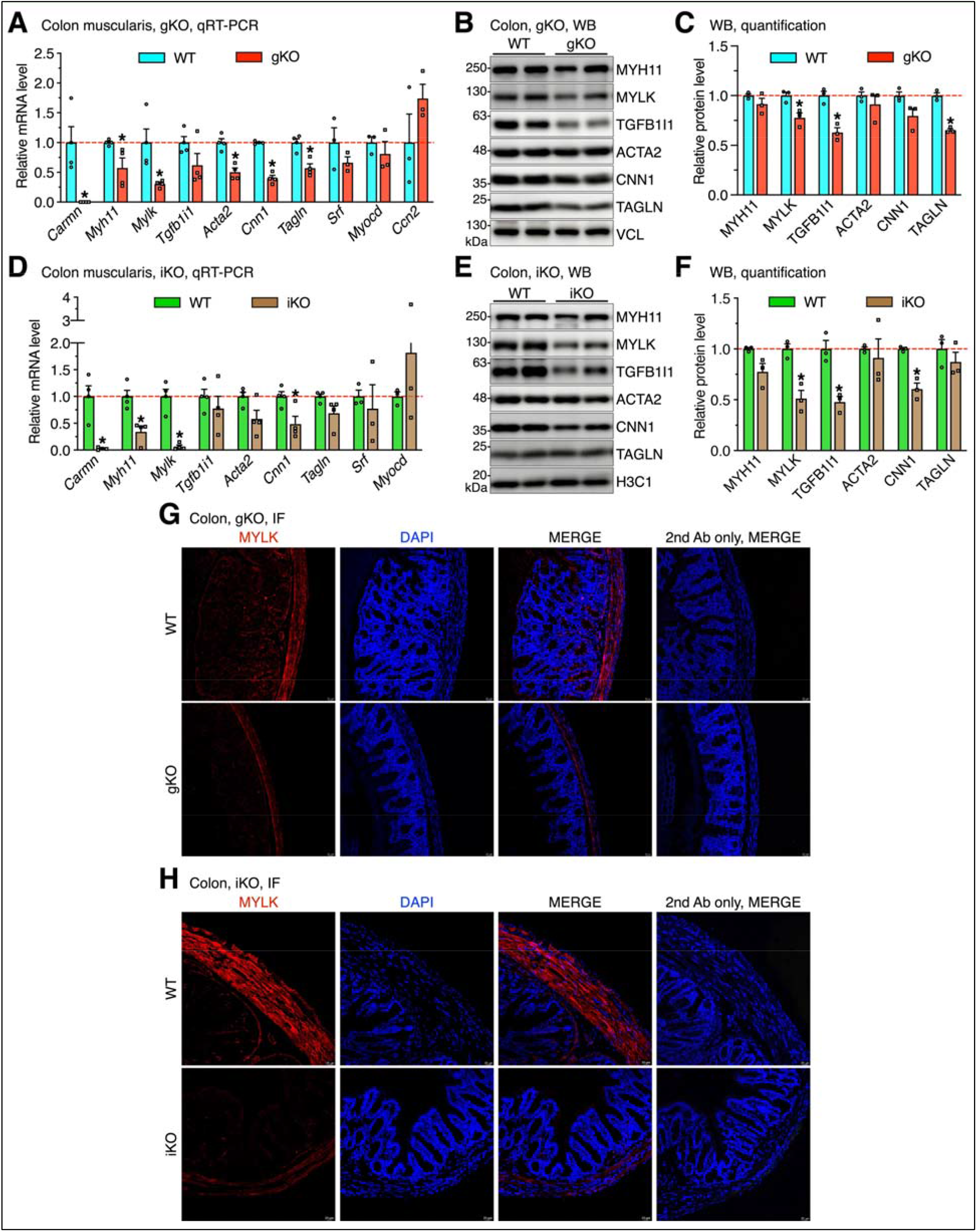
*Carmn* deficiency attenuates the expression of genes critical for SMC contraction. (**A-C**) Colon muscularis were isolated from the WT and *Carmn* gKO mice for (**A**) qRT-PCR or (**B**) Western blotting analysis. The expression level of mRNA (**A**) or protein (**C**) were quantified and presented relative to WT control (set to 1). N = 3-4; *p < 0.05; Unpaired Student *t* test. (**D-F**) Colon muscularis were harvested from the WT and *Carmn* iKO mice for (**D**) qRT-PCR and (**E**) Western blotting analysis. The expression level of (**D**) mRNA or (**F**) protein were quantified and presented relative to WT control (set to 1). N = 3-4; *p < 0.05; Unpaired Student *t* test. (**G-H**) Colon sections prepared from (**G**) *Carmn* gKO or (**H**) iKO mice and their respective control mice were stained with anti-MYLK antibody (red). Nuclei were counterstained with DAPI (blue). Sections only stained with second antibody served as the negative control.

### *CARMN* is critical for contraction of human colonic SMCs by regulating *MYLK* gene expression

We next sought to explore the translational relevance of *CARMN*-mediated contractile function in humans by using human colonic smooth muscle cells (HuCoSMCs). To examine whether *CARMN* regulates contractile gene expression in human GI SMCs, HuCoSMCs were transfected with control or *CARMN* antisense oligonucleotide (ASO) for 48 hours. qRT-PCR and Western blot analysis revealed that silencing of *CARMN* significantly increases expression of the proliferative gene PCNA and the matrix gene *CCN2* while attenuating the expression of almost all contractile genes examined except *MYH11* at both the mRNA and protein levels, compared to silencing control cells (**Figure 8A-C**). Since MYLK-mediated phosphorylation of RLC is prerequisite for the initiation of SMC contraction, we examined KCl and Cch-mediated MYLK-pMLC signaling pathway by Western blotting in HuCoSMCs with or without *CARMN*. The results from these experiments consistently showed that knocking down *CARMN* effectively attenuates the MYLK expression, thereby blocking MYLK-mediated phosphorylation of RLC (pMLC) (**Figure 8D-E**). To determine whether depletion of *CARMN* causes a functional loss of contractile competence of SMCs, we performed collagen gel contraction assays using HuCoSMCs in the presence or absence of *CARMN*. Results from this assay showed an approximate 50% decrease in contractile activity in *CARMN-*depleted HuCoSMCs (**Figure 8F-G**). It has been shown that the binding of the transcriptional complex of MYOCD and SRF to CArG elements residing in the promoter regions of SMC contractile genes is an important mechanism for regulating SM-specific gene expression, thereby maintaining the SMC contractile phenotype ^**51**^. Moreover, our recent study showed that *Carmn* directly binds with MYOCD, but not SRF to potentiate MYOCD-mediated myogenic function ^**26**^. Since *MYLK* has been known as a SRF target gene ^52^ and its SMC-specific expression overlaps with *CARMN* (**Figure 1D**), we hypothesize that *MYLK* is a novel target of *CARMN*/MYOCD/SRF complex in visceral SMCs. To test this hypothesis, we performed luciferase assays using WT or CArG mutant of *Mylk* gene reporter with or without co-transfection of expression plasmids of *Carmn* and *Myocd*. We also performed a similar luciferase assay in *Lmod1* gene reporter because we found depletion of *CARMN* leads to down-regulation of *LMOD1* in HuCoSMCs (**Figure 8A-C**) and *LMOD1* has been reported as an SRF/MYOCD-dependent gene and its mutation causes megacystis microcolon intestinal hypoperistalsis syndrome (MMIHS) in mouse and human ^**6**^. Data from these luciferase reporter assays in HuCoSMCs demonstrated that *Carmn* alone has no effect on the basal activity of *Mylk* and *Lmod1* gene promoters but significantly enhances MYOCD-mediated transactivation activity in a CArG-dependent manner (**Figure 8H**). Taken together, these results suggest that *Carmn* is indispensable for visceral SMC contraction by maintaining cell-cell connectivity and regulating SRF/MYOCD complex-dependent *MYLK* and *LMOD1* gene expression and MYLK-dependent contractile signaling *in trans*. Global deletion or SMC-specific deletion of *Carmn* in mice results in attenuated expression of *MYLK* and comprised cell-cell connectivity, leading to premature lethality due to the gut dysmotility-induced pseudo-obstruction (**Figure 8I**).

**Figure 8.**
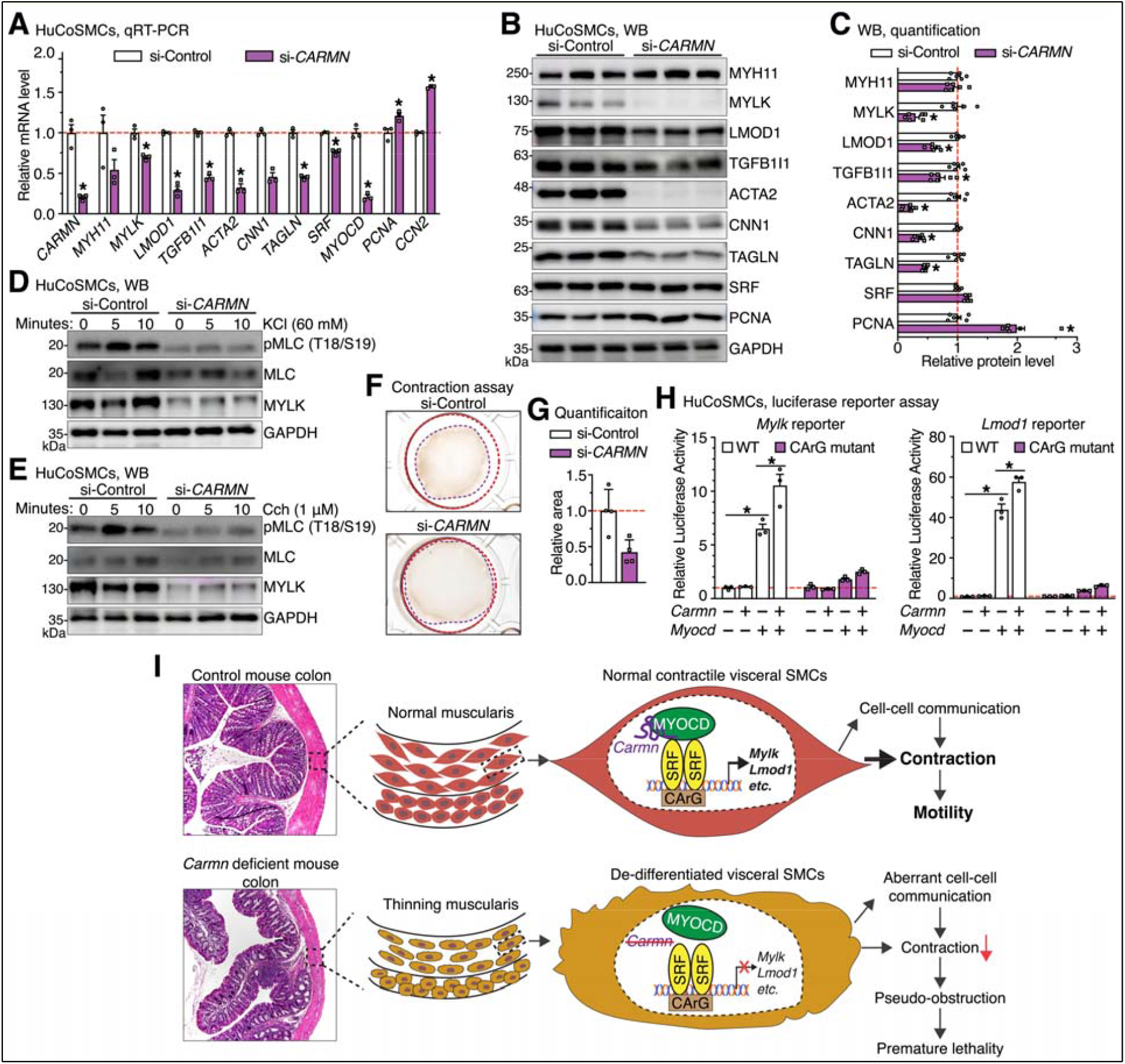
Depletion of *CARMN* impairs the contractile phenotype of human colonic SMCs. (**A**) *CARMN* or control phosphorothioate-modified ASO were transduced into human colonic SMCs (HuCoSMCs) for 48 hours to deplete the endogenous *CARMN* expression. Total RNA was then harvested for qRT-PCR analysis. The relative mRNA levels were quantified and presented relative to silencing control cells (set to 1). N = 3; *p < 0.05; Unpaired Student *t* test. **(B)** *CARMN* or control phosphorothioate-modified ASO were transduced into human colonic SMCs for 72 hours and then total protein was harvested for Western blotting analysis. (**C**) Densitometric analysis of relative protein level as shown in “**B**”. N = 6; *p < 0.05; Unpaired Student *t* test. (**D-E**) *CARMN* or control phosphorothioate-modified ASO were transduced into human colonic SMCs for 72 hours and cells were then treated with (**D**) 60 mM KCl or (**E**) the muscarinic agonist Carbachol (Cch, 1 µM) for the indicated time. Total protein was harvested for Western blotting analysis. (**F**) human colonic SMCs were transfected with control or *CARMN* phosphorothioate-modified ASO for 48 hours and then seeded onto 24-well plate for collagen contractility assay. (**G**) Quantitative analysis of the collagen contractility as shown in “**F**”. N = 4; *p < 0.05; unpaired Student *t* test. (**H**) Luciferase reporter assays were performed to examine the effects of *Carmn* on MYOCD-induced promoter activity of SMC-specific genes *Lmod1* and *Mylk* in which harbor wild-type (WT) or mutated CArG box. N = 3; *p < 0.05; 2-way analysis of variance. (**I**). Schematic diagram summarizing the major findings of this study.

## Discussion

CIPO is a rare but life-threatening disease characterized by severe intestinal dysmotility. Functional and histopathologic studies have shown that GI dysmotility in CIPO patients results from multiple mechanisms, including defects in neurons, SMCs or interstitial cells of Cajal (ICC) ^**53**^. Mutations affecting the expression and function of proteins of the contractile apparatus and cytoskeleton are the most common cause of myopathic CIPO ^**3**^. While lncRNAs have important epigenetic roles in SMC, their role in CIPO is poorly understood. Our study revealed a surprisingly critical role of a SMC-specific lncRNA, *CARMN*, in regulating visceral SMC contractility. Our findings raise the possibility that polymorphisms in the *CARMN* gene could lead to loss of expression and/or function, which possibly could contribute to some cases of myopathic CIPO.

Herein, we demonstrated that loss of *Carmn* expression in mice critically impairs colonic motility, disrupts colonic homeostasis and eventually results in lethal colonic pseudo-obstruction. To the best of our knowledge, our study is the first to show that a lncRNA is essential for maintaining visceral SMC contractility. The impact of loss of *Carmn* is seen with global deletion prior to development and SMC-specific inducible ablation in adulthood, with both mouse models developing a severe GI motility disorder with degenerative visceral myopathy, resembling SM dysfunction-mediated CIPO in humans. Future studies are warranted to examine whether there is a reduction in expression level or loss-of-function of mutations of *CARMN* in patients with CIPO. Furthermore, our comprehensive snRNA-seq analysis revealed that *Carmn* deficiency affects neurexins (NRXNs) and Neuregulins (NRGs)-mediated neuronal signaling. NRXNs and NRGs are synaptic cell-adhesion molecules that connect pre- and postsynaptic neurons at synapses, mediate trans-synaptic signaling, and shape neural network properties by specifying synaptic functions ^54^. How *Carmn* expression in SMC affects NRXN and NRG signaling remains undefined. Moreover, SNP mutations within NRXN are associated with susceptibility to Hirschsprung’s disease ^55^. We propose that *CARMN* may play a broader role, not only in SMCs per se, but also in regulating cell-cell communications that are crucial for coordinating GI motility.

We also found that pregnant *Carmn* heterozygous mice develop a dystocia phenotype, likely due to impaired uterine contractions. Notably, starting from day 21 when the pups were weaned to regular chow diet from milk feeding, 60% of *Carmn* global knockout mice succumbed to the diet switch within 2 weeks, suggesting that *Carmn* is required for force overload response in both uterine and GI tract SMCs. A limitation of our study is that we were unable to assess the specific functional role of *Carmn* in postnatal uterine SMCs because the inducible *Myh11*-CreER^T2^ transgene we used is only active in male mice ^**30**^. A new SMC-specific Cre mouse model in which Cre is active in both sexes of mice is required to address this question.

During the preparation of this study, Vacante et al reported on the use of CRISPR/Cas9-mediated genome editing to constitutively knockout *Carmn* in mice ^**27**^. Using this mouse model, they found that loss of *Carmn* reduced the expression of SMC contractile genes in the mouse aorta, similar to our findings in the mouse GI tract. However, this study did not report a gastrointestinal phenotype in *Carmn* KO mice. Recently, Ni et al used antisense GapmeRs specific to *Carmn* to knockdown *Carmn* in mice ^**28**^. They showed that despite the reduced efficacy of antisense GapmeRs, achieving approximately 50% efficiency in knocking down *Carmn*, they significantly decreased expression of SM-specific contractile genes in the mouse aorta. However, this study also did not report any GI phenotype in mice treated with *Carmn* antisense GapmeRs. The reasons for these apparent discrepancies are not clear although we can speculate that they may relate to a focus on the vasculature rather than the GI tract, residual or altered function of the *Carmn* in the CRISPR knockout, differences in the genetic background of mice and/or the low efficiency of knockdown *in vivo* by GapmeRs.

Previous studies have shown that both the SRF transcription factor and its cofactor, MYOCD are necessary for the development and maintenance of the visceral SMC contractile phenotype. Targeted deletion of either gene in SMCs of adult mice resulted in severe gut dysmotility, characterized by weak peristalsis and dilation of the digestive tract ^**11**, **12**^. Notably, the SMC-specific inducible *Carmn* KO mice in our study exhibited remarkable similarities in the gross, microscopic and molecular findings of the SM-specific inducible *Myocd* and *Srf* deletion ^**10**, **11 12, 56**^. This is not completely surprising given that *Carmn* was found to bind to and potentiate MYOCD/SRF complex function as we previously reported in vascular SMCs ^**26**^, but the extent of regulation by *Carmn* is remarkable. Our findings support a role for *CARMN* as a critical binding partner of the MYOCD/SRF complex that has a vitally important role in maintaining the function of visceral SMCs. However, the lethal GI myopathy phenotype resulting from *Myh11* promoter-driven Cre has precluded us from assessing a potential role of *Carmn* in vascular SMCs *in vivo*. To address this important question, future studies will need to be conducted by using a vascular SMC-specific Cre mouse model.

Using bulk RNA-seq and snRNA-seq approaches to investigate mechanisms, we identified that *Carmn* deficiency disrupts the homeostasis of the colon and impairs colonic motility mainly through down-regulation of an array of SMC contractile genes especially the *Mylk* gene. Consistent with previous studies showing that *MYLK* is an SRF/MYOCD target gene ^**52**^, we found that *Carmn*-induced transcriptional activation of the *MYLK* gene promoter is CArG-dependent. It is well accepted that SMC regulatory light chain (RLC) phosphorylation by calcium/calmodulin-dependent MLCK is a prerequisite for SM contraction and critical for maintaining the physiological movement of hollow organs ^38, 57-59^. We found that deletion of *Carmn* in both murine visceral SMCs *in vivo* and human colonic SMCs *in vitro* not only significantly down-regulated *Mylk* gene expression, but also attenuated KCl and Cch-induced RLC phosphorylation. Together with the previous study showing that inducible SMC-specific deletion of *Mylk* in mouse results in markedly reduced RLC phosphorylation, gut motility and premature lethality, our data suggests *MYLK*, as a *CARMN* target gene, at least in part, is responsible for mediating *CARMN* function in visceral SMCs. However, we cannot rule out the possibility that other down-regulated genes such as *Dmpk* and *Chrm2* may play additional functional roles since these genes are also SRF/MYOCD targets genes and have been shown to be important for smooth muscle contraction ^56, 60^. Interestingly, a recent computational study suggested that *Carmn* could regulate the expression of downstream genes including *MYLK* via formation of RNA-DNA-DNA triplexes ^61^. Although this idea is provocative, confirmation on whether *Carmn* regulates the expression of SMC-contractile genes via RNA-DNA interactions needs to be tested in wet lab experiments. Based on data showing a reduction in a novel population of neuron-like SMCs in *Carmn* KO mice, our snRNA-seq data further suggest that *Carmn* regulates cellular connectivity between SMCs and other cell types. We propose that cell-cell decoupling may partly account for the decreased contractility observed in the GI tracts of *Carmn* KO mice. Collectively, our data show that SM-specific lncRNA *Carmn* plays a critical role in maintaining visceral SMC homeostasis that is likely to occur through multiple mechanisms.

*CARMN* is an evolutionarily conserved smooth muscle cell-enriched lncRNA that was initially identified as the host gene for *MIR143/145* cluster ^**26**^, which is the best-characterized microRNA pair regulating SMC differentiation and phenotypic modulation ^**62-65**^. Using bulk RNA-seq data we found that *Carmn* deletion robustly decreases the expression of *miR143/145* cluster in colon tissues (**Supplemental Table 2**). However, previous studies demonstrated that germline deletion of the *miR143/145* cluster in mice resulted in no overt developmental defects at baseline, although it did contribute to a defect in epithelial regeneration in response to intestinal injury ^**66**^. In the current study, we found that both global deletion and SMC-specific deletion of *Carmn* resulted in a lethal colonic pseudo-obstruction phenotype, which is very different from the negligible phenotype observed in mice absent the *miR143/145* cluster. Together, these results indicate that the ability of *Carmn* to maintain colonic SMC homeostasis is independent of the *miR143/145* cluster although *Carmn* deletion can affect *miR143/145* expression. This finding is also consistent with recent studies showing that *CARMN* can regulate the plasticity of vascular SMC phenotypes independent of the *MIR143/145* cluster in vitro ^**26-28**^. However, *miR145* was reported to strongly express in zebrafish gut smooth muscle and loss of *miR145* in zebrafish resulted in deficits of smooth muscle maturation and contraction ^**67**, **68**^, a similar finding to our mouse study. Given that the putative mouse *Carmn* ortholog was not annotated and identified in zebrafish until in our recent report ^**26**^ and that the *Carmn* gene is localized within the *miR143/145* gene locus, we speculate that the GI phenotype observed in zebrafish with knockout of *miR145* is likely due to an accidental deletion of *Carmn*. To confirm this, a *Carmn*-specific knockout/knockdown zebrafish model should be generated to directly address the distinct roles of *Carmn* versus *miR145*.

In summary, our study has uncovered a novel role of the lncRNA, *CARMN*, as a major regulator of gastrointestinal motility and viability through actions in visceral SMCs that serve to maintain contractile function and intercellular connectivity. A greater understanding of the GI roles of lncRNAs such as *CARMN* adds to our tool kit in helping to understand and diagnose human visceral myopathy diseases such as CIPO.

## Supporting information

Supplemental Material including supplemental methods, tables and figures

## What You Need to Know

### Background and Context

Visceral smooth muscle cells (SMCs) are critical for regulating motility of gastrointestinal (GI) tract. Loss of GI motility causes several motility diseases including chronic intestinal pseudo-obstruction (CIPO) in humans. The functional role of long non-coding RNAs (lncRNAs) in regulating GI motility is largely unexplored.

### New Findings

*Carmn* is a SMC-specific lncRNA that is conserved in humans and mice. *Carmn* deficiency in mice results in premature lethality due to colonic pseudo-obstruction by compromising intestinal motility through reduced expression of contractile genes and impairing cell-cell connectivity in the colonic muscularis.

### Limitations

Study limitations include having not yet validated the sub-population of neuron-like SMCs in mice and lack of evidence to show the decreased expression of *CARMN* in human CIPO patients’ samples.

### Impact

*Carmn* is indispensable for maintaining GI SMC contractile function in mice, but also it suggests that loss of function of *CARMN* may contribute to human visceral myopathy such as CIPO.

