## Supplemental Material including supplemental methods, tables and figures for "The LncRNA *Carmn* is a Critical Regulator for Gastrointestinal Smooth Muscle Contractile Function and Motility"

[#] **Corresponding author**

**This supplementary information includes:**

**Part I: Supplemental text**

Supplemental Material and Methods, and Supplemental Reference

**Part II: Supplemental Tables**

**Table S1:** Primers used for genotyping of *Carmn* KO/GFP KI mouse and SMC lineage tracing mouse

**Table S2**: Bulk RNA-seq data sheets

**Table S3**: SnRNA-seq data sheets

**Table S4**: Antibodies used for Western blotting and immunofluorescence staining

**Table S5:** Primers used for qRT-PCR

**Part III: Supplemental Figures**

**Figure S1:** Marker genes used to identify the cell types in human and mouse GI tissues and visualization of *Carmn* in jejunum tissues of *Carmn* GFP KI mouse

**Figure S2:** Global deletion of Carmn in mice causes jejunum distension and SMC degeneration

**Figure S3**: Inducible deletion of *Carmn* in mice causes jejunum distension and SMC degeneration

**Figure S4:** *Carmn* deficiency in mice impairs the colonic contractility

**Figure S5**: Bulk RNA-seq analysis on the jejunum and colon muscularis of *Carmn* WT and iKO mice

**Figure S6:** SnRNA-seq analysis on the colon muscularis of *Carmn* WT and iKO mice

**Figure S7:** Cell-cell communication analysis on the colon muscularis of *Carmn* WT and iKO mice

**Figure S8**: *Carmn* deficiency decreases the expression of *Mylk* in jejunum muscularis of *Carmn* gKO and iKO mice.

**Figure S9**: Uncropped Western blots for Figure 7

**Figure S10**: Uncropped Western blots for Figure 8

### Supplemental Material and Methods

#### Bioinformatics analysis of public bulk RNA-seq and scRNA-seq datasets

For bulk RNA-seq analysis, we downloaded the bulk RNA-seq dataset of human colon and small intestine from the GTEx Portal database [1], and analyzed the expression of top lncRNAs in the human tissues.

To identify the most abundant lncRNAs in human colon tissues at the single cell level, we re-analyzed the single-cell transcriptome dataset for 62,849 cells isolated from 6-10 weeks post-conception developing human gut including duojejunum, ileum and colon (https://www.gutcellatlas.org) [2]. The downloaded count matrix of scRNA-seq data were analyzed using the DESC package [3]. Genes expressed in fewer than 3 cells, and cells expressing fewer than 200 or more than 6000 genes, or containing >15% mitochondrial genes were removed. Count matrix of scRNA-seq datasets for adult human and mouse colon was downloaded from Gene Expression Omnibus (GEO) database (GSE156905) [4]. The obtained count matrix was processed with R packages "Seurat" for quality control, dimensionality reduction and clustering [5]. Genes expressed in fewer than 3 cells, and cells expressing fewer than 150 or more than 6000 genes, or containing >20% mitochondrial genes were removed. Uniform Manifold Approximation and Projection (UMAP) plots were generated for visualization of different cell clusters.

#### Sectioning, Hematoxylin/Eosin (HE) and immunofluorescence (IF) staining

Colon and Jejunum tissues were collected from the sex- and age-matched WT and KO mice, then fixed with 4% paraformaldehyde overnight. The fixed tissues were then embedded in paraffin, sectioned for HE staining or IF assay following standard protocols as previously described [6-8]. For directly visualizing GFP and IF in sections, colon and jejunum tissues were fixed with 4% PFA overnight, washed with 3 times with PBS, then kept in 30% sucrose overnight. Fixed tissues were embedded in optimal cutting temperature compound (OCT) and saved at -80$^{o}$C till cryo-sectioning. Cryo-sections were air-dried for 15 minutes and washed with PBS for 3 times. Sections were blocked and permeabilized with goat serum (10%, Thermo Fisher) plus 0.1% Tween-20 for 30 minutes, then incubated with MYH11 or MYLK antibody overnight at 4$^{o}$C. After washing with PBS, sections were incubated with secondary antibody diluted in the blocking buffer for 1 hour at room temperature. Following 3-time wash with PBS, sections were mounted with ProLong Gold anti-fade reagent with DAPI (Thermo Fisher) and imaged using a confocal microscopy (LSM 780 upright, Zeiss). The antibodies used were listed in **Supplemental Table 4**.

**Whole-gut transit time measurement**

To assess the mouse GI function, mice were fasted overnight and next day mice were fed with the artificial food (10% charcoal and 5% gum arabic mix in the water). Mice were then placed individually in cages, and the time taken for excretion of the first charcoal-stained feces was recorded as the whole-gut transit time.

**Wire myography**

The multiwire myography system (DMT 620M) was used for mounting proximal colon to measure the mechanical force. The system connects with vacuum and gassing units. A heating unit is provided in the myography interface, allowing for all chambers to be evenly maintained under physiological setting (37°C, and bubbled with 21% $O_2$ : 5% $CO_2$). The myograph system is also connected to a computer via an amplifier (Power Lab, AD Instruments) for continuous measurement of isometric tension using the data acquisition software (Lab Chart). Mouse colons were isolated from the control and gKO mice at postnatal day 20 or from the control and *Carmn* iKO mice at day 30 post the first tamoxifen injection. The internal contents and connective tissues were quickly removed in oxygenated Kreb's buffer. For isometric tension recordings, approximately 5 mm muscle strips with mucosa were cut from proximal colons and suspended in the circular direction in 5 mL organ bath containing oxygenated Kreb's buffer at 37°C. The tissues were allowed to equilibrate for 30 minutes before stimulation was applied and the basal tension was set to 1 gram. After equilibration, KCl (60 mM) or Carbachol (1 $\mu$M) were applied to elicit muscle contraction. The tension trace was recorded, and the contractility force was expressed as the peak amplitude during stimulation.

**Cell culture for human colonic smooth muscle cells (HuCoSMCs)**

HuCoSMCs (ScienCell Research Laboratories; Cat. #: 2940) and smooth muscle cell culture medium (ScienCell Research Laboratories; Medium, Cat. #: 1101; DPBS, Cat. #: 0303; Trypsin/EDTA, Cat. #: 0183; Trypsin Neutralizing Solution, Cat. #: 0113; PLL, Cat. #: 0403) were purchased from ScienCell Research Laboratories. HuCoSMCs were cultured in a humidified atmosphere of 95% air and 5% $CO_2$ at 37°C following the protocol provided by vendor. Sub-confluent HuCoSMCs (passages 3-5) were seeded in 6-well plate for Western blot or qRT-PCR. For cell contraction assay, HuCoSMCs were seeded onto 24-well plate.

**Antisense oligonucleotides (ASO) transfection**

Three phosphorothiote modified ASOs targeting the common region of human *CARMN* transcripts (*CARMN*-ASO-1, 5'-TCTGTGAAAGGTGATG-3'; *CARMN*-ASO-2, 5'-CAGAGTTCTTGCTTCTCTGA-3'; *CARMN*-ASO-3, 5'-AGGTTCCACTTCTTAACGAG-3') and scrambled ASO (5'-TCATACTATATGACAG-3') which was used as control were synthesized by Exiqon. Delivery of ASO (mixed *CARMN*-ASO-1, 2 and 3, or scrambled ASO) into primary HuCoSMCs were carried out by using Neon transfection system (Thermo Fisher) essentially following the manufacturer's protocol and was described in our previous report [6]. After transfection, HuCoSMCs were plated in 6-well plates for qRT-PCR or Western blot analysis, as indicated in the figure legends. For cell contraction assays, equal numbers of HuCoSMCs were plated in 24-well plates.

**Quantitative reverse transcription-PCR (qRT-PCR) analysis**

Total RNA was harvested from *Carmn* KO (iKO or gKO) and control mouse colon or jejunum muscularis tissues, or from HuCoSMCs with TRIzol reagent (Thermo Fisher, Cat. #: 15596018).

0.8 μg of RNA was utilized as template for RT with random hexamer primers using the High Capacity RNA-to-cDNA Kit (Thermo Fisher, Cat. #: 4374966). qRT-PCR was performed with respective gene-specific primers as listed in **Supplemental Table 5**. All samples were amplified in duplicate and all experiments were repeated at least 3 independent times. Relative gene expression was converted using the $2^{-\triangle\triangle CT}$ method against the internal control house-keeping gene *Gapdh* where $\triangle\triangle CT = (CT_{\text{experimental gene}} - CT_{\text{experimental } Gapdh}) - (CT_{\text{control gene}} - CT_{\text{control } Gapdh})$. The relative gene expression in control group is set to 1.

**Protein extraction and Western blotting**

Total protein lysates were extracted from mouse colon or jejunum muscularis tissues, or from HuCoSMCs by RIPA buffer (Fisher Scientific) plus 1% protease/phosphatase inhibitor cocktail (Fisher Scientific). After sonication and centrifugation of the samples, protein in the supernatant was quantified by BCA assay (Fisher Scientific) and resolved on a 7.5%,10% or 12.5% SDS-PAGE gel at 10 μg per lane as appropriate. Primary antibodies used in this study were listed in **Supplemental Table 4**. Secondary antibodies conjugated with horseradish peroxidase were then used to visualize the target proteins on blot. Image was acquired by Amersham ImageQuant 800 biomolecular imager (GE) and band densities were quantified using the Image J software.

**Cell contraction assay**

Forty-eight hours after transfection with scrambled ASO or mixed *CARMN* ASO, HuCoSMCs were trypsinized and used for a collagen contractility assay essentially following the manufacturer's protocol (Cell Biolabs, Cat. #: CBA-201).

**Bulk RNA-sequencing**

Colon or jejunum muscularis were isolated from *Carmn* iKO and control mice (day 30 post the 1st tamoxifen injection) for total RNA extraction with TRIzol reagent (Thermo Fisher, Cat. #: 15596018). The total RNA was then subjected to the whole transcriptome RNA-seq analysis for differential gene expression analysis (Washington University in St. Louis). Total RNA integrity was determined using the Agilent Bioanalyzer. Library preparation was performed with 1 μg of total RNA. Ribosomal RNA was removed by an RNase-H method using RiboErase kit (Kapa Biosystems). mRNA was then fragmented in reverse transcriptase buffer and heating to 94 degrees for 8 minutes. mRNA was reverse transcribed to yield cDNA using random hexamers and SuperScript III RT enzyme (Life Technologies) per manufacturer's instructions. A second strand reaction was performed to yield ds-cDNA. cDNA was blunt ended, had an A base added to the 3' ends, and then had Illumina sequencing adapters ligated to the ends. Ligated fragments were then amplified for 12 cycles using primers incorporating unique dual index tags. Fragments were sequenced on an Illumina NovaSeq-6000 using paired end reads extending 150 bases. RNA-seq reads were then mapped to mouse reference genome mm10 using STAR version 2.0.4b [9]. The raw count for genes annotated in Ensembl GRCm38.76 were calculated with Subread:featureCount version 1.4.5 [10] and counts per million mapped reads (CPM) was obtained to quantify relative expression of transcripts. Differential expression analysis was performed with the R package edgeR [11]. For each comparison, only those genes with average CPM > 5 in at least one group were analyzed and genes with fold changes (FC) larger than 1.5 and p-value adjusted (padj) less than 0.05 were considered statistically significant. Volcano plots were generated using R program. GO (Gene Ontology) analysis was carried out by g:Profilter [12].

**Nuclei isolation and FACS sorting for single nucleus RNA-sequencing (snRNA-seq)**

Snap-frozen colon muscularis tissues isolated from *Carmn* iKO and control mice (at day 30 post the 1$^{st}$ tamoxifen injection) were thawed on ice and cut into small pieces. Crude nuclei were isolated by homogenization following the protocol provided by 10X Genomics (CG000375). Isolated nuclei were resuspended in PBS +1% BSA (Miltenyi Biotec, Cat. #: 130-091-376) + 1U/μl RNase inhibitor (Roche, Cat. #: 03335402001) supplemented with 1% 7AAD (Sigma, Cat #: SML1633). Nuclei were then sorted using BD FACSAria II cell sorter. Nuclei were isolated from debris by gating on the 7AAD positive signal. Doublets were removed with gating on an FSC-A vs FSC-H. The sorted nuclei were then saved in the solution of 1% BSA with 1U/μl RNase inhibitor and were adjusted concentration by spinning and resuspending the pellet in chilled nuclei buffer provided 10X Genomics.

**snRNA-seq**

Nuclei isolated from the colon muscularis of *Carmn* iKO and control mice as described above were processed for snRNA-seq. For each sample, sorted nuclei were counted using Countless II FL automated cell counter following trypan blue staining (Thermo Fisher, Cat. #: T10282) and subsequently processed according to the 10X Chromium single cell 3' protocol. Completed 10X libraries were sequenced using an Illumina NextSeq-500/550 high output kit v2 at the Georgia Cancer Center-Integrated Genomics and Bioinformatics Shared Resources at Augusta University. The Cell Ranger Single Cell software suite 3.0 (10X Genomics) using STAR aligner [9] was employed to align sequencing reads to the mouse pre-mRNA reference genome (mm10) and to generate BAM files as well as filter gene-barcode matrices on the High Performance Cluster. The cellranger count function was used to create filtered gene-cell expression matrices by removing cell barcodes not represented in nuclei.

**Pre-processing and data analysis workflow**

The count matrix of snRNA-seq dataset generated from *Carmn* iKO and control samples were processed with R packages Seurat [13] for quality control, dimensionality reduction and clustering. Genes expressed in fewer than 3 cells, and cells expressing fewer than 150 or more that 6000 genes, or containing > 20% mitochondrial genes were removed. Subsequently, *Carmn* iKO and control Seurat objects were integrated into one Seurat object containing 5,072 nuclei in the primary analysis dataset using FindIntegrationAnchors functions and IntegrateData functions. Principal component (PC) analysis was performed using RunPCA. The number of useful PCs was determined using the ElbowPlot function and graph-based clustering was performed with the FindClusters function at a resolution of 0.5. UMAP plots were generated for visualization of different cell clusters. FindAllMarkers created the list of genes differentially expressed in each cluster compared to all other nuclei based on the Willcoxon rank-sum test and employing a cutoff of a minimal absolute log fold change difference of 0.25 and expression in minimal 10% of the nuclei in the respective cluster. The re-clustering analysis of SMCs employed the subset function to create a new Seurat object from the primary analysis dataset with subsequent graph-based clustering with the FindClusters function at a resolution of 0.5. Signaling pathway enrichment analysis in the SMC clusters were carried out by g:Profilter [12]. The SMCs trajectories were analyzed using the Monocel 2 R package [14]. Cell-cell interaction was evaluated using the CellChat R package [15].

The transcriptomics data including both bulk and snRNA-seq datasets generated in this study have been deposited in the Sequences Read Archive at the NCBI under accession # GSE199157.

**Luciferase reporter assay**

All promoter genes were constructed by cloning fragments of promoters into the pGL2B luciferase reporter vector (Promega, Cat. #: E1960). The WT or CArG-mutant promoter of *Lmod1* gene has been reported by the previous publication [16]. The WT or CArG-mutant promoter of *Mylk* gene was synthesized by GenScript Biotech based on the previous publication [17]. The sequence of luciferase reporter vectors carrying the WT or CArG-mutant promoter of *Lmod1* and *Mylk* genes were sequenced by Genewiz. Transfection into HuCoSMCs was carried out with X-tremeGENE 9 DNA transfection reagent (Roche, Cat. #: 06365787001) following the protocol provided by the manufacturer. The level of promoter activity was evaluated by measurement of the firefly luciferase relative to the internal control Renilla luciferase using the dual luciferase assay system essentially as described by the manufacturer (Promega). A minimum of three independent transfections were performed.

**Supplemental Table 1**

**Primers used for genotyping of *Carmn* KO/GFP KI mouse and SMC-lineage tracing mouse.**

F: Forward; R: reverse

| Primer Name | Sequence (5'-3') |
|---|---|
| *Carmn* P1 | F: CACTTCCCACTTCACCCCACCTT |
| *Carmn* P2 | R: CCCTGCTGTCCATTCCTTATTCC |
| *Carmn* P3 | R: GTGCCTCAGTTTCCCTACCTATCTTA |
| *Myh11*-CreER$^{T2}$ | F: TCCAACCTGCTGACTGTG |
| *Myh11*-CreER$^{T2}$ | R: TCAGAGTTCTCCATCAGGG |
| mTmG Common | F: CTCTGCTGCCTCCTGGCTTCT |
| mTmG WT | R: CGAGGCGGATCACAAGCAATA |
| mTmG Mutant | R: TCAATGGGCGGGGGTCGTT |

**Supplemental Table 4**

**Antibodies used for Western blotting (WB) and immunofluorescence (IF) staining**

| Protein | Vendor | Catalog | Dilution for WB | Dilution for IF |
|---|---|---|---|---|
| MYH11 | Abcam | Ab224804 | | 1: 300 |
| MYLK | Abcam | Ab76092 | 1: 1000 (for human cells) | 1: 300 |
| GFP | Invitrogen | A11120 | | 1: 200 |
| MYH11 | Biomedical Technologies | BT-562 | 1: 5000 (for mouse tissue)<br>1: 1000 (for human cells) | |
| MYLK | Sigma | M7905 | 1: 5000 (for mouse tissue) | |
| LMOD1 | Proteintech | 15117-1-AP | 1: 5000 | |
| TGFB1I1 | BD | 611164 | 1: 5000 | |
| ACTA2 | Abcam | Ab5694 | 1: 10000 (for mouse tissue)<br>1: 5000 (for human cells) | |
| CNN1 | Sigma | C2687 | 1: 5000 (for mouse tissue) | |
| CNN1 | Abcam | Ab46794 | 1: 2000 (for human cells) | |
| TAGLN | Abcam | Ab10135 | 1: 2000 | |
| pMLC(T18/S19) | CST | 3674S | 1: 1000 | |
| MLC | CST | 8505S | 1: 1000 | |
| SRF | CST | 5147S | 1: 1000 | |
| PCNA | Santa Cruz | SC-56 | 1: 1000 | |
| VCL | Sigma | V4505 | 1: 5000 | |
| GAPDH | Santa Cruz | SC-47724 | 1: 1000 | |
| H3C1 | CST | 4499S | 1: 1000 | |

**Supplemental Table 5**

**Primers used for qRT-PCR** (F: Forward; R: Reverse)**.**

| Gene name | Species | Sequence (5'-3') |
|---|---|---|
| *Carmn* | Mouse | F: CAACAGCCGCCCAGCTCCCAGAACT |
| | | R: TCCCATGTCTTCAGAAGATGTCTCC |
| *Myh11* | Mouse | F: CACAGGAAACTTCGCAGTGA |
| | | R: TTCTGTTTTCCCTGACATGGT |
| *Mylk* | Mouse | F: TGCATGGTGGTGAATGGGTCAGGGA |
| | | R: GTCACCTCCTTCCTGATGTCTGGAC |
| *Tgfb1i1* | Mouse | F: GCCTCTGTGGCTCCTGCAATAAAC |
| | | R: CTTCTCGAAGAAGCTGCTGCCTC |
| *Acta2* | Mouse | F: ATGCTCCCAGGGCTGTTTTCCCAT |
| | | R: GTGGTGCCAGATCTTTTCCATGTCG |
| *Cnn1* | Mouse | F: TCATCTGCACCTCTGCTTTG |
| | | R: GGGCCAGCTTGTTCTTTACT |
| *Tagln* | Mouse | F: TGACATGTTCCAGACTGTTGACCTCT |
| | | R: CTTCATAAACCAGTTGGGATCTCCAC |
| *Srf* | Mouse | F: GATGGAGTTCATCGACAACAAGCTG |
| | | R: CCCTGTCAGCGTGGACAGCTCATA |
| *Myocd* | Mouse | F: GTTCAGCTACCCTGGGATGCACCAA |
| | | R: GGCCTGGTTTGAGAGAAGAAACACC |
| *Gapdh* | Mouse | F: GGCATTGCTCTCAATGACAA |
| | | R: TGTGAGGGAGATGCTCAGTG |
| *CARMN* | Human | F: AATTAGTTGAGAAGCAGTGACACC |
| | | R: CAGAGTTCTTGCTTCTCTGACATC |
| *MYH11* | Human | F: GGTCACGGTTGGGAAAGATGA |
| | | R: GGGCAGGTGTTTATAGGGGTT |

| | | |
|---|---|---|
| *MYLK* | Human | F: CCCGAGGTTGTCTGGTTCAAA |
| | | R: GCAGGTGTACTTGGCATCGT |
| *LMOD1* | Human | F: GAGGCCATGCTCAACTTCTG |
| | | R: CTCTCCATTCTTGGCATCTG |
| *TGFB1I1* | Human | F: GCCTCTGTGGCTCCTGCAATAAAC |
| | | R: CTTCTCGAAGAAGCTGCTGCCTC |
| *ACTA2* | Human | F: ATGCTCCCAGGGCTGTTTTCCCAT |
| | | R: GTGGTGCCAGATCTTTTCCATGTCG |
| *CNN1* | Human | F: CTGTCAGCCGAGGTTAAGAAC |
| | | R: GAGGCCGTCCATGAAGTTGTT |
| *TAGLN* | Human | F: TGACATGTTCCAGACTGTTGACCTCT |
| | | R: CTTCATAAACCAGTTGGGATCTCCAC |
| *SRF* | Human | F: GATGGAGTTCATCGACAACAAGCTG |
| | | R: CCCTGTCAGCGTGGACAGCTCATA |
| *MYOCD* | Human | F: ATTCAGCTACCTAGGGATGCACCAAG |
| | | R: GGCCTGGTTTGAAAGAAGAGACACC |
| *PCNA* | Human | F: AGGTGTTGGAGGCACTCAAG |
| | | R: CATTGCCGGCGCATTTTAGT |
| *GAPDH* | Human | F: TTGGTATCGTGGAAGGACTC |
| | | R: ACAGTCTTCTGGGTGGCAGT |

### Supplemental Figures and Legends

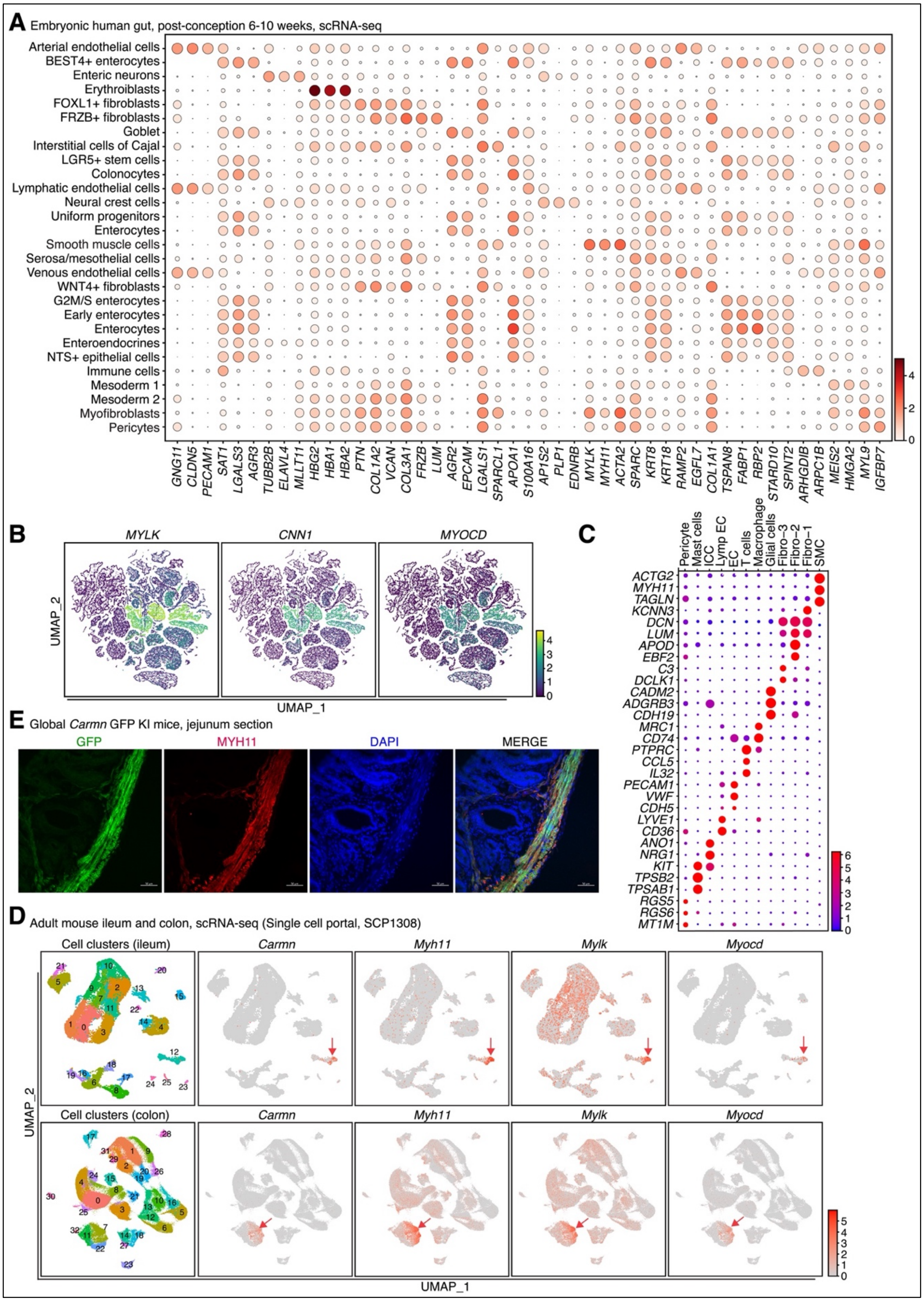

**Figure S1. Marker genes used to identify the cell types in human and mouse GI tissues and visualization of *Carmn* in jejunum tissues of *Carmn* GFP KI mouse**. (**A**) Dot plot showing the marker genes used to identify the cell types of embryonic human gut tissue. (**B**) UMAP showing the cell types and the expression of SMC-specific genes *MYLK*, *CCN1* and *MYOCD* in the cell types of embryonic human gut as revealed by the scRNA-seq. (**C**) Dot plot showing the marker genes used to identify the cell types of adult human colon. (**D**) UMAP to visualize the expression of *Carmn* and SMC-specific genes *Myh11*, *Mylk* and *Myocd* in adult mouse ileum (upper panel) and colon (bottom panel). Arrows point to *Carmn* positive cell clusters. (**E**) Direct visualization of GFP and immunostaining of MYH11 to examine the specific cellular localization of GFP signal in jejunum tissue dissected from *Carmn* K/W (Het) mice. Scale bar: 20 μm.

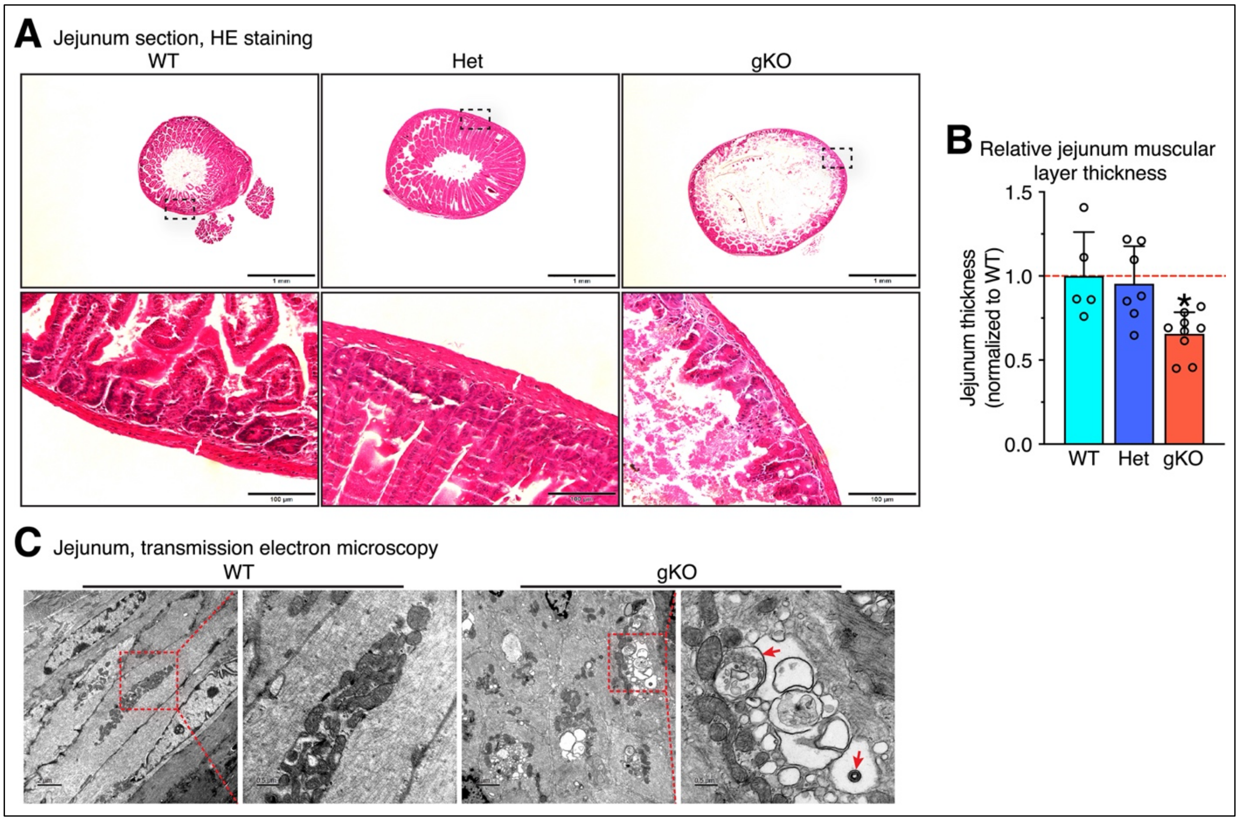


**Figure S2. Global deletion of *Carmn* in mice causes jejunum distension and SMC degeneration.** (**A**) Hematoxylin and eosin (HE) staining on the transverse sections of jejunum from WT, Het and *Carmn* gKO mice. The boxed area is magnified underneath each respective panel. Scale bar: 1 mm and 100 μm in the upper and bottom panel, respectively. (**B**) The thicknesses of jejunum muscularis layers were measured and plotted. N = 5-9; *$p < 0.05$; Unpaired Student *t* test. (**C**) Representative transmission electron microscopy (TEM) images demonstrate mitochondrial and endoplasmic reticulum ultrastructure of WT and *Carmn* gKO jejunum SMCs. The boxed area is magnified on the right. Arrows denote the representative laminated autophagic vesicles.

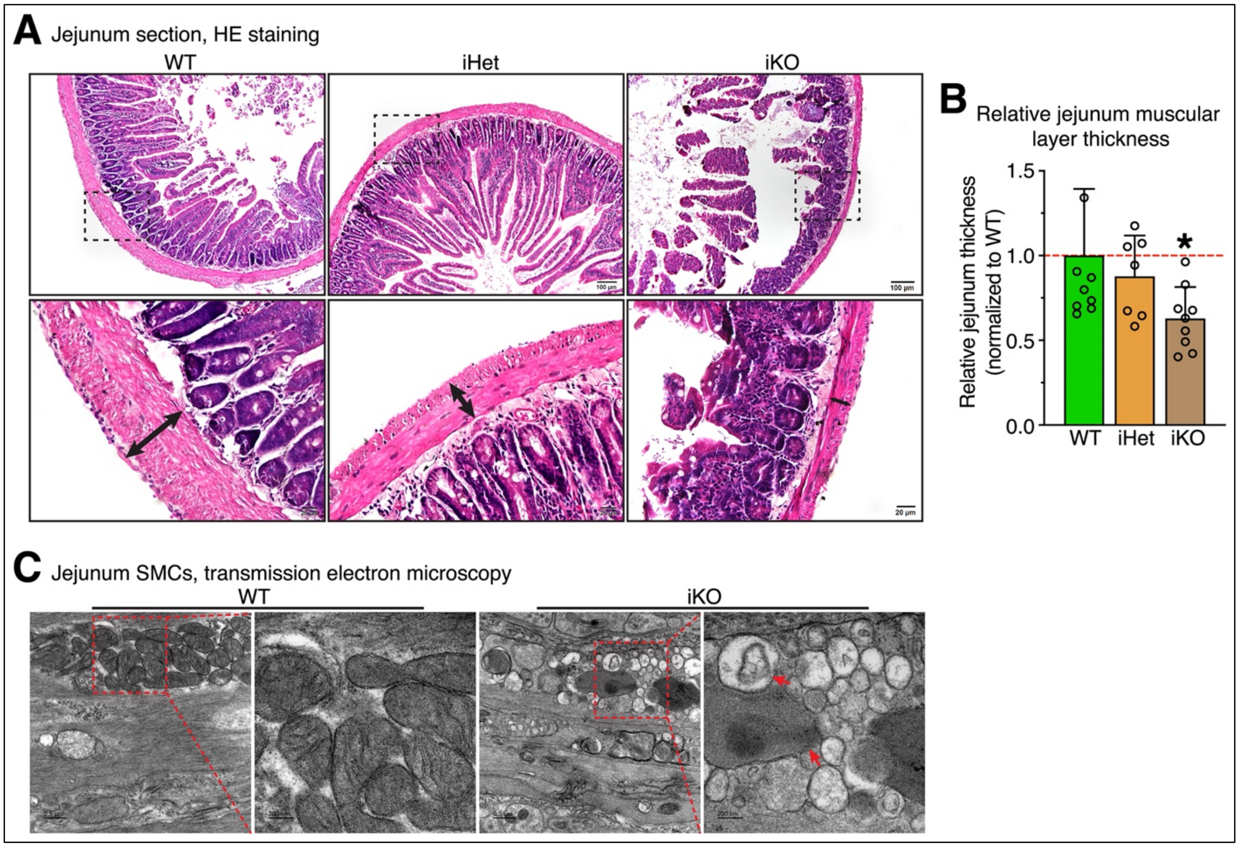


**Figure S3. Inducible deletion of *Carmn* in adult mice causes jejunum distension and SMC degeneration.** (**A**) Hematoxylin and eosin (HE) staining on transverse sections of jejunum isolated from WT, iHet and *Carmn* iKO mice. The boxed area is magnified on the bottom. The double-head arrows indicate the muscularis layer. Scale bar: 100 μm and 20 μm in the upper and bottom panel, respectively. (**B**) The thicknesses of muscularis layers were measured in jejunum transverse section and plotted relative to the WT. N = 7-9; *p < 0.05; Unpaired Student *t* test. (**C**) Representative transmission electron microscopy (TEM) images showing the mitochondrial morphology in WT and *Carmn* iKO jejunum SMCs. The boxed area is magnified to the right. Arrows point to the representative autophagic vesicles.

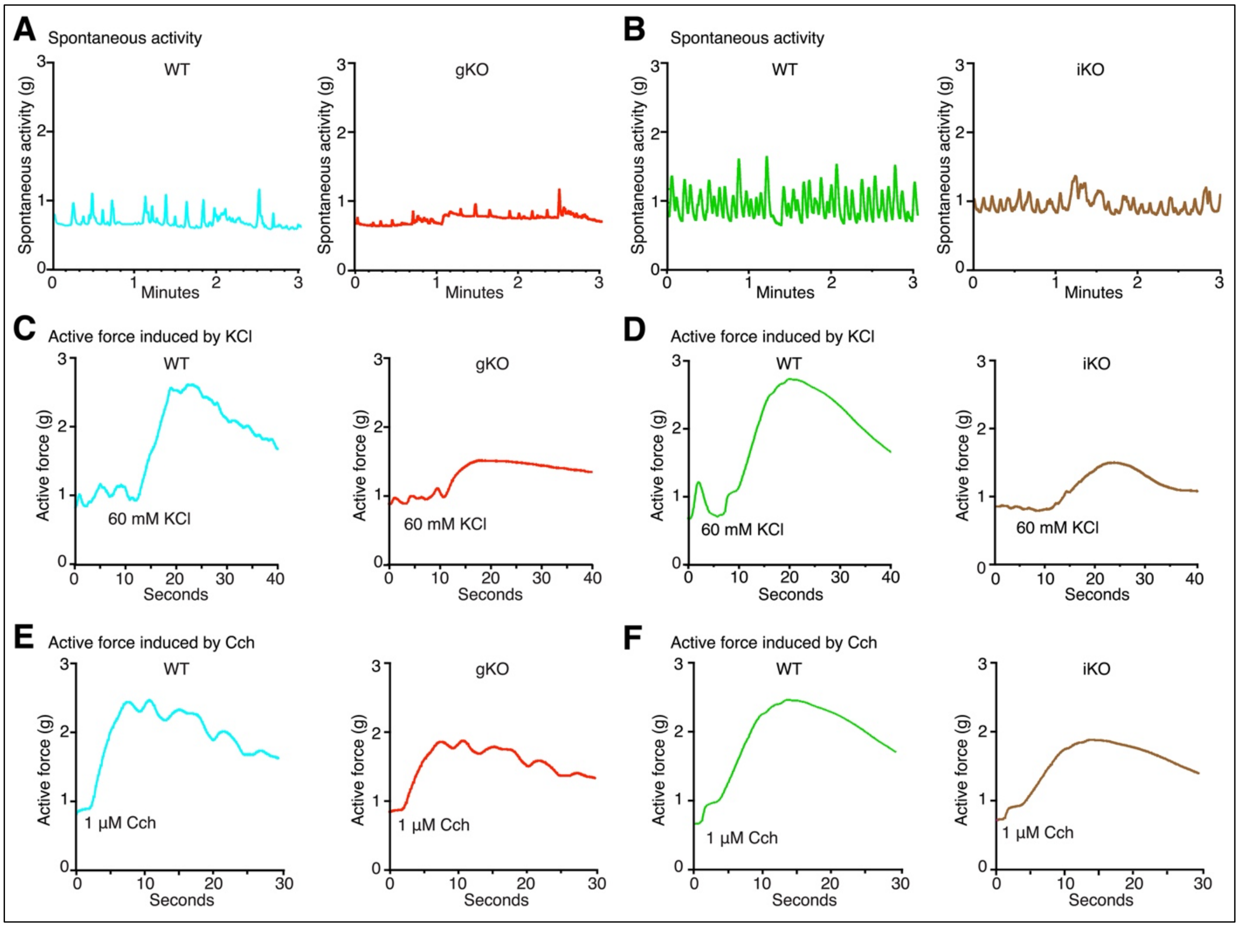


**Figure S4. *Carmn* deficiency in mice impairs the colonic contractility.** (**A**-**B**) Representative spontaneous contraction recording on colonic segments from (**A**) WT control and *Carmn* gKO or (**B**) iKO mice. (**C-D**) Representative contraction recordings induced by 60 mM KCl on colonic rings from (**C**) *Carmn* gKO or (**D**) iKO mice and their respective WT control mice. **(E-F)** Representative recordings of contraction induced by 1 µM Cch on colonic rings from (**E**) *Carmn* gKO or (**F**) iKO mice and their respective WT control mice. Superimposed contracting recordings from both control and KO mice are presented in Figure 4.

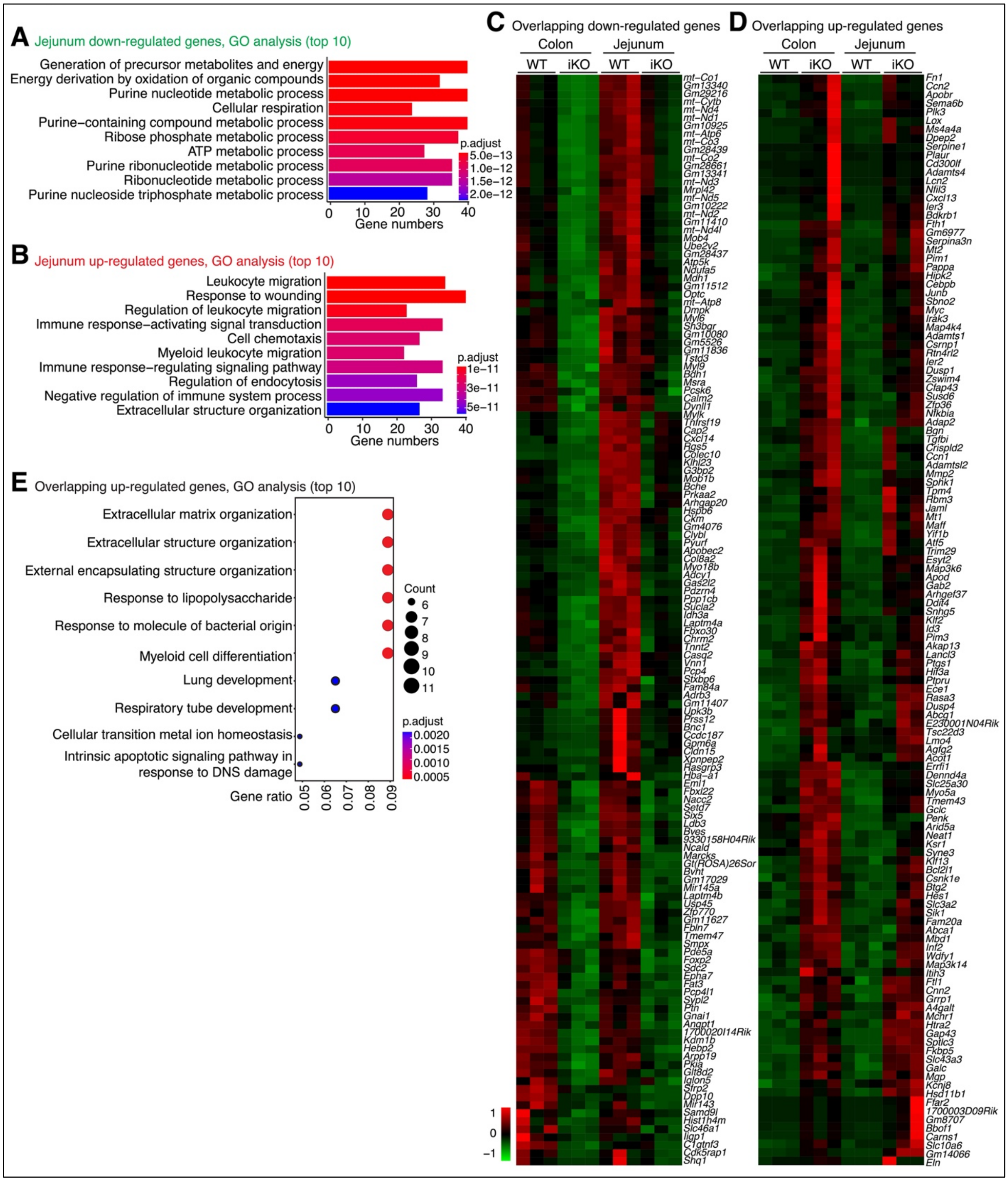


**Figure S5. Bulk RNA-seq analysis on the jejunum and colon muscularis of *Carmn* WT and iKO mice.** (**A-B**) GO analysis showing the top 10 enriched GO biological processes for (**A**) the 561 significantly down-regulated or (**B**) 635 up-regulated genes in *Carmn* iKO jejunum muscularis. (**C-D**) Heatmaps showing the expression change of overlapping (**C**) down- or (**D**) up-regulated genes in both colon and jejunum of *Carmn* iKO mice, respectively. (**E**) GO analysis showing the top 10 enriched GO biological processes for the 128 overlapping up-regulated genes in both colon and jejunum muscularis of *Carmn* iKO mice.

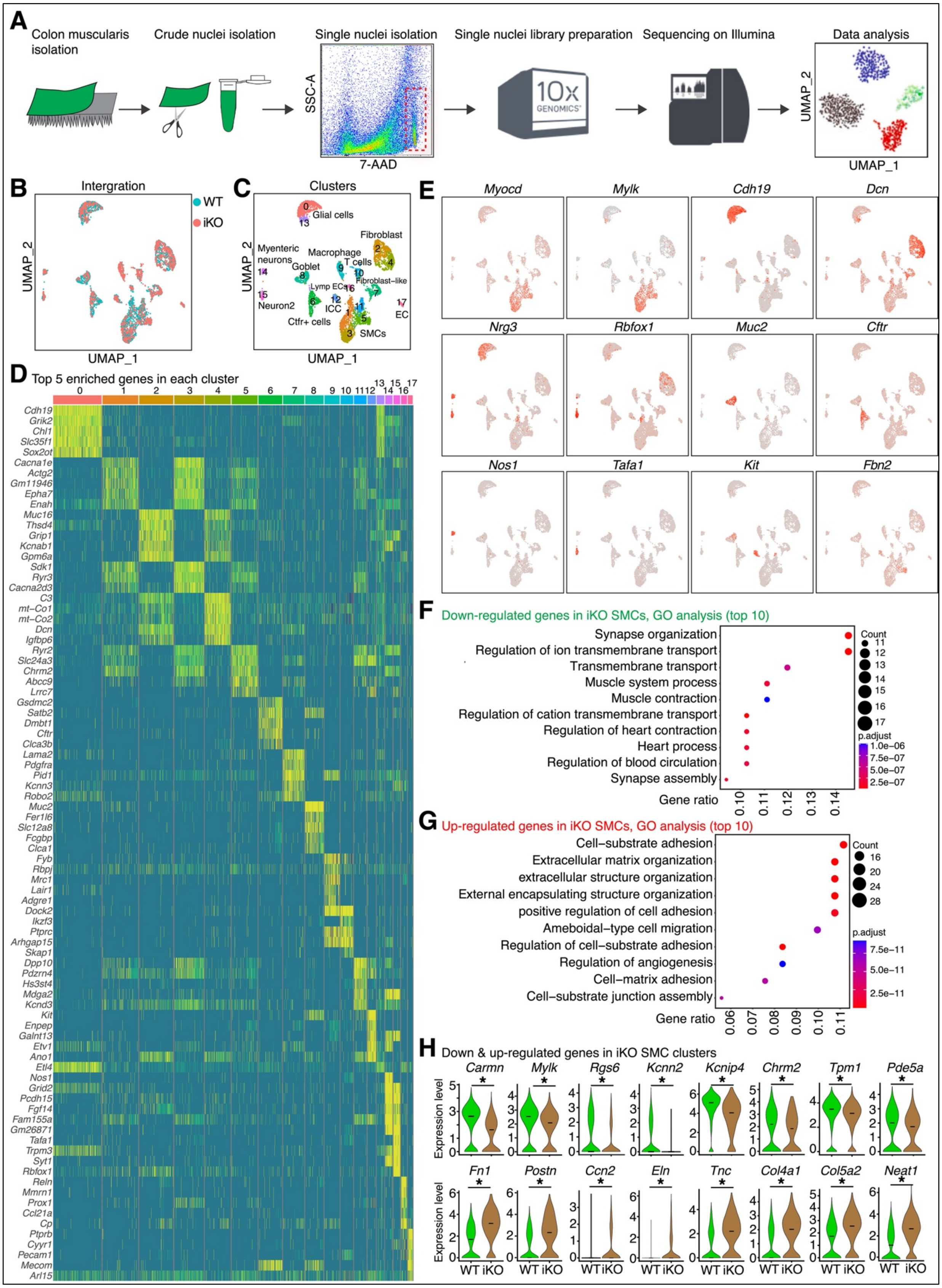
A
Colon muscularis isolation
Crude nuclei isolation
Single nuclei isolation
Single nuclei library preparation
Sequencing on Illumina
Data analysis
SSC-A
7-AAD
UMAP_2
UMAP_1
B
Intergration
WT
iKO
C
Clusters
Glial cells
Fibroblast
Myenteric neurons
Macrophage
T cells
Goblet
Fibroblast-like
Lymp ECs
Neuron2
ICC
EC
Cftr+ cells
SMCs
D
Top 5 enriched genes in each cluster
E
F
Down-regulated genes in iKO SMCs, GO analysis (top 10)
Synapse organization
Regulation of ion transmembrane transport
Transmembrane transport
Muscle system process
Muscle contraction
Regulation of cation transmembrane transport
Regulation of heart contraction
Heart process
Regulation of blood circulation
Synapse assembly
Gene ratio
Count
p.adjust
G
Up-regulated genes in iKO SMCs, GO analysis (top 10)
Cell-substrate adhesion
Extracellular matrix organization
extracellular structure organization
External encapsulating structure organization
positive regulation of cell adhesion
Ameboidal-type cell migration
Regulation of cell-substrate adhesion
Regulation of angiogenesis
Cell-matrix adhesion
Cell-substrate junction assembly
Gene ratio
Count
p.adjust
H
Down & up-regulated genes in iKO SMC clusters
Expression level
WT iKO

**Figure S6. SnRNA-seq analysis on the colon muscularis of *Carmn* WT and iKO mice.** (**A**) Workflow for the procedures of snRNA-seq analysis on the colon muscularis of *Carmn* iKO and WT mice. (**B**) UMAP showing the integration of cell clusters identified by snRNA-seq on the colon muscularis of *Carmn* iKO and WT mice. (**C**) UMAP visualization of the cell clusters and types based on the gene expression profile in the individual cell. (**D**) Heatmap showing the top 5 enriched genes in each cell cluster in comparison to the other cell clusters. (**E**) UMAP visualization of a subset of selected genes that are used to define the cell types. (**F-G**) The top 10 most enriched GO biological processes for the (**F**) down- and (**G**) up-regulated genes between *Carmn* iKO and WT SMCs as revealed by snRNA-seq. **(H)** Violin plots showing the significantly down- (upper panel) or up-regulated expression (bottom panel) of a subset of representative genes in the colonic SMCs of *Carmn* iKO compared to that of WT mice. *FDR $< 0.05$; Differential analysis with Seurat package.

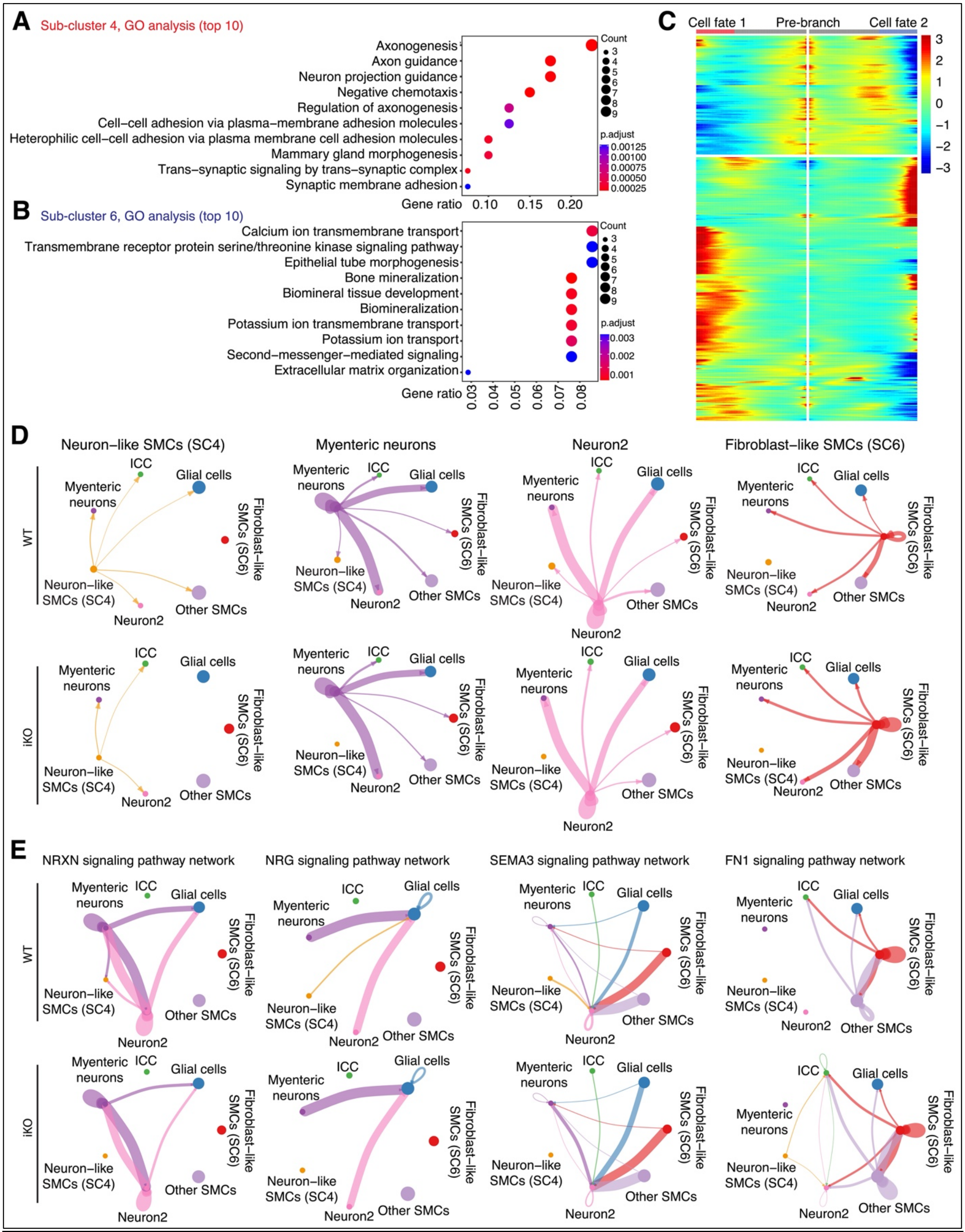


**Figure S7. Cell-cell communication analysis on the colon muscularis of *Carmn* WT and iKO mice.** (**A-B**) The top 10 most enriched GO biological processes for genes enriched in (**A**) SMC sub-cluster 4 (neuron-like SMCs) and (**B**) 6 (fibroblast-like SMCs). (**C**) Heatmap showing the pseudotime trajectory analysis of SMC fate. (**D**) Comparison of the significantly changed outgoing cell-cell communications between WT (top panels) and *Carmn* iKO (bottom panels) mouse colon muscularis. (**E**) Comparison of NRXN, NRG, SEMA3 and FN1-mediated signaling among the cell types identified in mouse colon muscularis of WT (top panels) and *Carmn* iKO (bottom panels) mice.

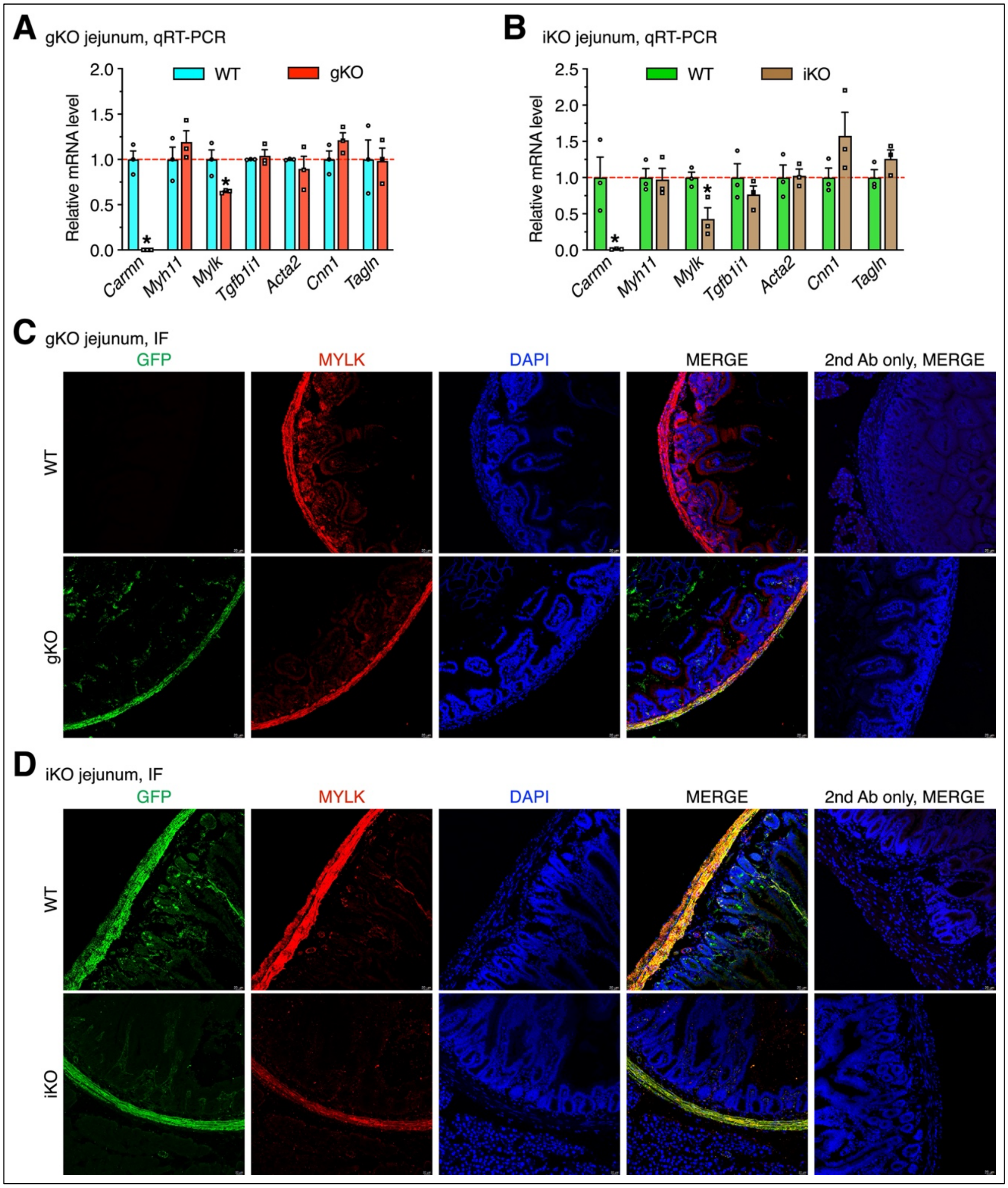


**Figure S8. *Carmn* deficiency decreases the expression of *Mylk* in jejunum muscularis of *Carmn* gKO and iKO mice.** (**A-B**) Jejunum muscularis were isolated from the (**A**) *Carmn* gKO or (**B**) iKO mice and their respective WT control mice for qRT-PCR. The relative mRNA levels were quantified and presented relative to WT control (set to 1). N = 3; *$p < 0.05$; Two-way ANOVA. (**C-D**) Jejunum sections prepared from (**C**) *Carmn* gKO or (**D**) iKO and their respective WT control mice were stained with anti-GFP (green) and anti-MYLK (red) antibody. Nuclei were counterstained with DAPI (blue). Sections only stained with second antibody served as the negative control.

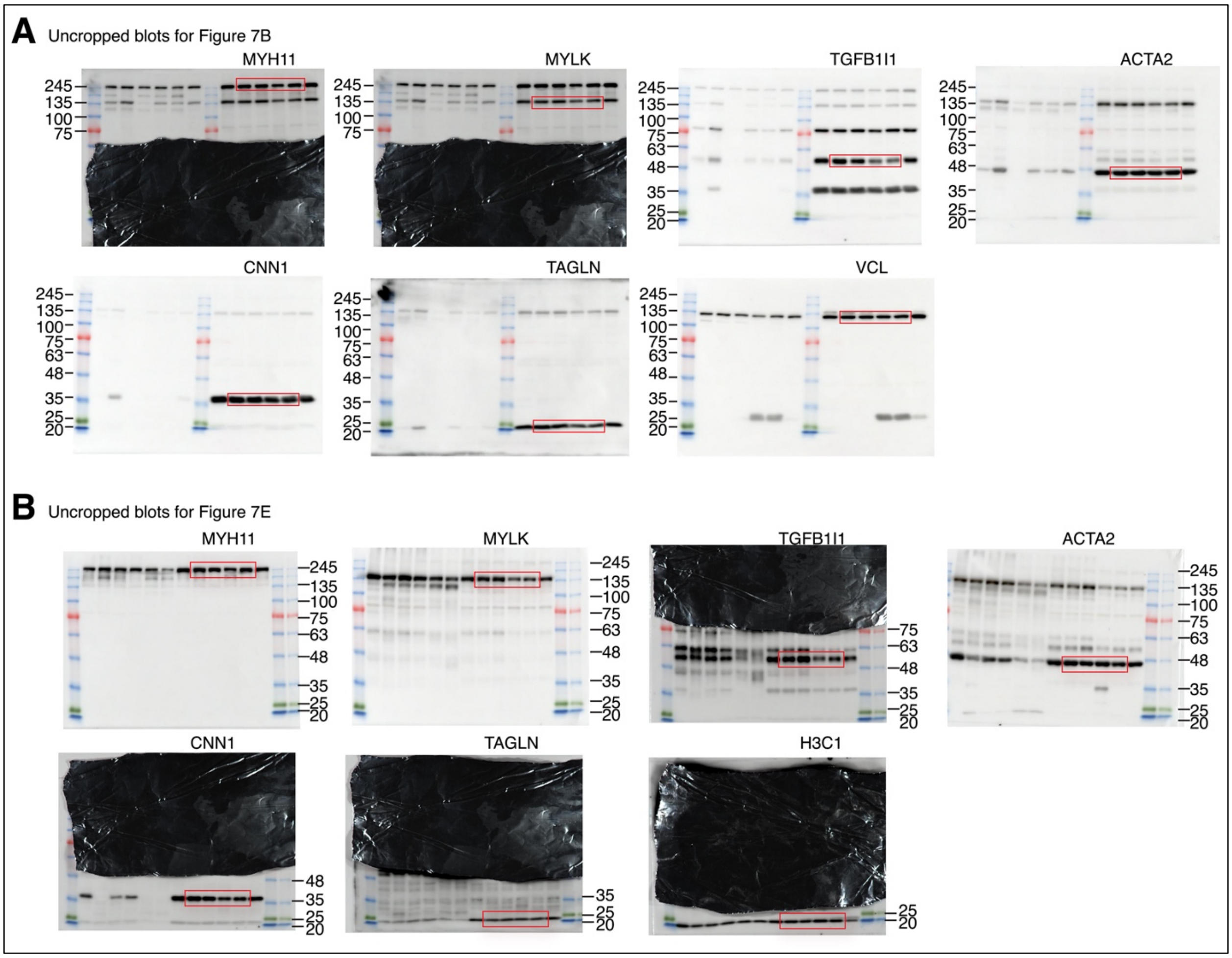


**Figure S9. Uncropped Western blots for Figure 7.** Uncropped original Western blots for (**A**) Figure 7B and (**B**) Figure 7E, respectively.

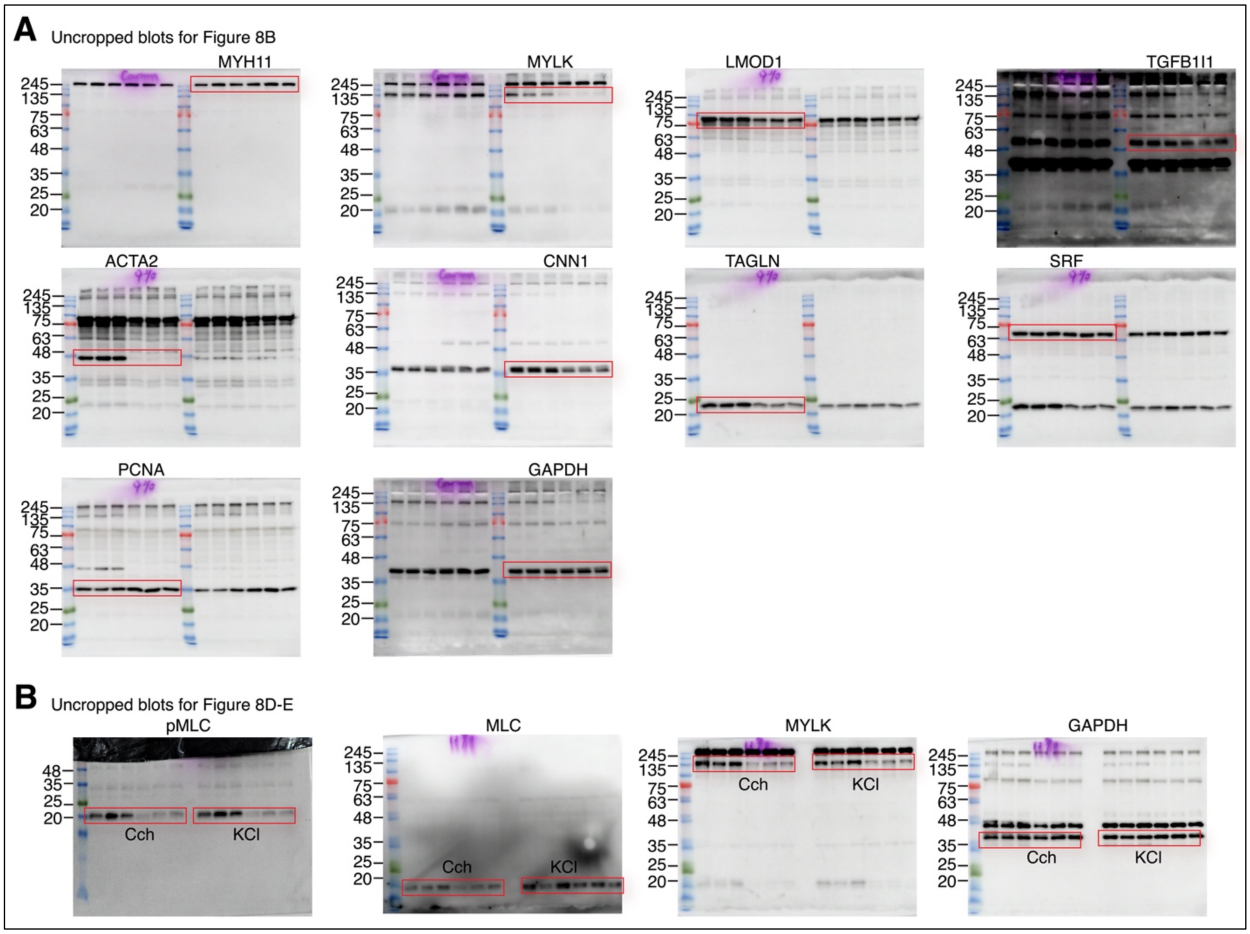


**Figure S10. Uncropped Western blots for Figure 8.** Uncropped original Western blots for (**A**) Figure 8B and (**B**) Figure 8D-E, respectively.
